# GraphGPSM: a global scoring model for protein structure using graph neural networks

**DOI:** 10.1101/2023.01.17.524382

**Authors:** Guangxing He, Jun Liu, Dong Liu, Zhang Guijun

## Abstract

The scoring models used for protein structure modeling and ranking are mainly divided into unified field and protein-specific scoring functions. Although protein structure prediction has made tremendous progress since CASP14, the modeling accuracy still cannot meet the requirements to a certain extent. Especially, accurate modeling of multi-domain and orphan proteins remains a challenge. Therefore, an accurate and efficient protein scoring model should be developed urgently to guide the protein structure folding or ranking through deep learning. In this work, we propose a protein structure global scoring model based on equivariant graph neural network (EGNN), named GraphGPSM, to guide protein structure modeling and ranking. We construct an EGNN architecture, and a message passing mechanism is designed to update and transmit information between nodes and edges of the graph. Finally, the global score of the protein model is output through a multilayer perceptron. Residue-level ultrafast shape recognition is used to describe the relationship between residues and the overall structure topology, and distance and direction encoded by Gaussian radial basis functions are designed to represent the overall topology of the protein backbone. These two features are combined with Rosetta energy terms, backbone dihedral angles, and inter-residue distance and orientations to represent the protein model and embedded into the nodes and edges of the graph neural network. The experimental results on the CASP13, CASP14, and CAMEO test sets show that the scores of our developed GraphGPSM have a strong correlation with the TM-score of the models, which are significantly better than those of the unified field score function REF2015 and the state-of-the-art local lDDT-based scoring models ModFOLD8, ProQ3D, and DeepAccNet etc. The modeling experimental results on 484 test proteins demonstrate that GraphGPSM can greatly improve the modeling accuracy. GraphGPSM is further used to model 35 orphan proteins and 57 multi-domain proteins. The results show that the average TM-score of the models predicted by GraphGPSM is 13.2% and 7.1% higher than that of the models predicted by AlphaFold2. GraphGPSM also participates in CASP15 and achieves competitive performance in global accuracy estimation.

## 1 Introduction

Proteins participate in cell metabolism, maintain immune functions, and are essential substances for life activities. The importance of protein structure lies in the fact that it can make proteins have specific shapes and activities, so that they can perform certain biological functions [1]. Determining 3D protein structure can help us understand protein functions in cellular processes and modulate the functions through drug discovery [2]. While protein structure determination based on experiments remains time-consuming and expensive [3, 4], considerable protein structure prediction methods based on state-of-the-art deep learning techniques have been proposed to predict the 3D structure of proteins from sequences. Scoring models are an indispensable part of protein structure prediction and are used to guide protein structure modeling and select the best model. Breakthroughs have been made in protein structure prediction through artificial intelligence, effectively utilizing the power of multiple sequences of protein families [2], such as AlphaFold2 [5] and RoseTTAFold [6]. Nevertheless, the modeling of orphan and multi-domain proteins still needs to be improved substantially when effective MSA data are not available. In this case, a reliable scoring model must be designed to guide protein structure modeling.

Protein structure scoring models can roughly be divided into two categories: unified field and protein-specific scoring models. Unified field scoring models describe the general properties of proteins through the physical and chemical knowledge of proteins and statistical potentials from the PDB database [7, 8]. For example, van der Waals statistical potentials describe short-range attractive and repulsive forces as a function of atom-pair distance, hydrogen bond potentials are used to describe partial covalent bond interactions, and solvent potentials describe the hydrophilicity and hydrophobicity of residues [7]. These potentials are important for capturing the structural specificities that underlie protein folding, functions, and interactions. With the rapid development of deep learning-based protein structure prediction, protein-specific scoring models are widely exploited. The non-end-to-end versions of RoseTTAFold [6] and trRosetta [9] convert deep learning predictions of inter-residue distance and orientations into bound energy potentials that guide protein modeling along with uniform field energies. IPTDFold constructs distance energy potentials to perform protein topology adjustment and local dihedral angle optimization [10]. Uniform field scoring models reflect the universal properties of proteins, but they have low accuracy, are computationally complex, and lack the unique properties of different proteins. This deficiency can be compensated by protein-specific field energy constraint potentials. However, constraint potentials constructed on the basis of deep learning methods rely heavily on protein coevolutionary information, and it is difficult to guide modeling well in the absence of coevolutionary information.

Protein model quality assessment can also be regarded as a scoring model, which is used to select predicted protein models and provide key information for protein refinement [11–13]. Most model quality assessment methods mainly focus on the local lDDT, and the global score of the model is obtained by averaging the lDDT of all residues. Therefore, although lDDT-based scoring can capture the trend of accuracy changes at local residues, it does not reflect the overall folding accuracy of protein models well. The latest research found that for complex proteins, the metric constructed by the average interface plDDT [5] and the number of interface contacts can indeed describe the relationship between the complex protein interfaces [14]. On CASP15, our developed lDDT-based model quality assessment method DeepUMQA3 (group name: GuijunLab RocketX) achieved the best performance in the accuracy estimation of protein complex interface residues [37]. However, it performed poorly in the estimation of global folding accuracy. The reason is that the average plDDT [5] of the entire complex usually results in a poor-quality assessment, suggesting that both single chains in a complex are often predicted very well, but their relative orientation may still be incorrect [14]. For multi-domain proteins, we hypothesized that another global metric is needed to assess reasonably the accuracy of the global topology, rather than just averaging the local lDDT. Therefore, developing a global scoring model for proteins that is independent of coevolutionary information is necessary.

In this work, we design a protein global scoring model, GraphGPSM, based on equivariant graph neural network (EGNN). Atomic-level backbone features encoded by Gaussian radial basis functions, residue-level ultrafast shape recognition (USR), Rosetta energy terms, distance and orientations, one-hot encoding of sequences, and positional embedding of residues are used to characterize protein structures. These features are configured to the nodes and edges of the initial graph and combined with coordinate embedding to construct the initial architecture of the EGNN. A dense messaging network is formed by stacking multiple layers of this architecture. Finally, the score of the structure model is produced by a multilayer perceptron consisting of dropout layers, activation functions, and linear layers. Experiments reveal a good correlation between the score of GraphGPSM and the true TM-score of protein structure models, which is better than those of the existing models with global lDDT-based scoring. Furthermore, GraphGPSM can be used to guide structure modeling more accurately. Particularly, GraphGPSM can significantly improve the accuracy of models predicted by AlphaFold2 on our developed orphan and multi-domain protein test set. The performance of GraphGPSM (group name: GuijunLab Threader) on CASP15 also makes it into a top-ranked server for global accuracy prediction.

## 2 Materials and Methods

The pipeline of GraphGPSM is shown in Figure 1. Our method consists of four parts, which are data preparation, feature encoder, network architecture, and model training. For the protein structure to be scored, it is characterized by atomic-level backbone features encoded by Gaussian radial basis functions, residue-level USR, Rosetta energy terms, inter-residue distance and orientations, one-hot encoding of sequence, and sinusoidal position encoding of residues. Then, these features are embedded into the nodes and edges of the graph to form an initial graph. A complete graph propagation path is built by stacking multiple graphs as layers of the network and using a message passing mechanism between these layers. Finally, the score of the protein model is output through a multilayer perceptron.

**Figure 1.**
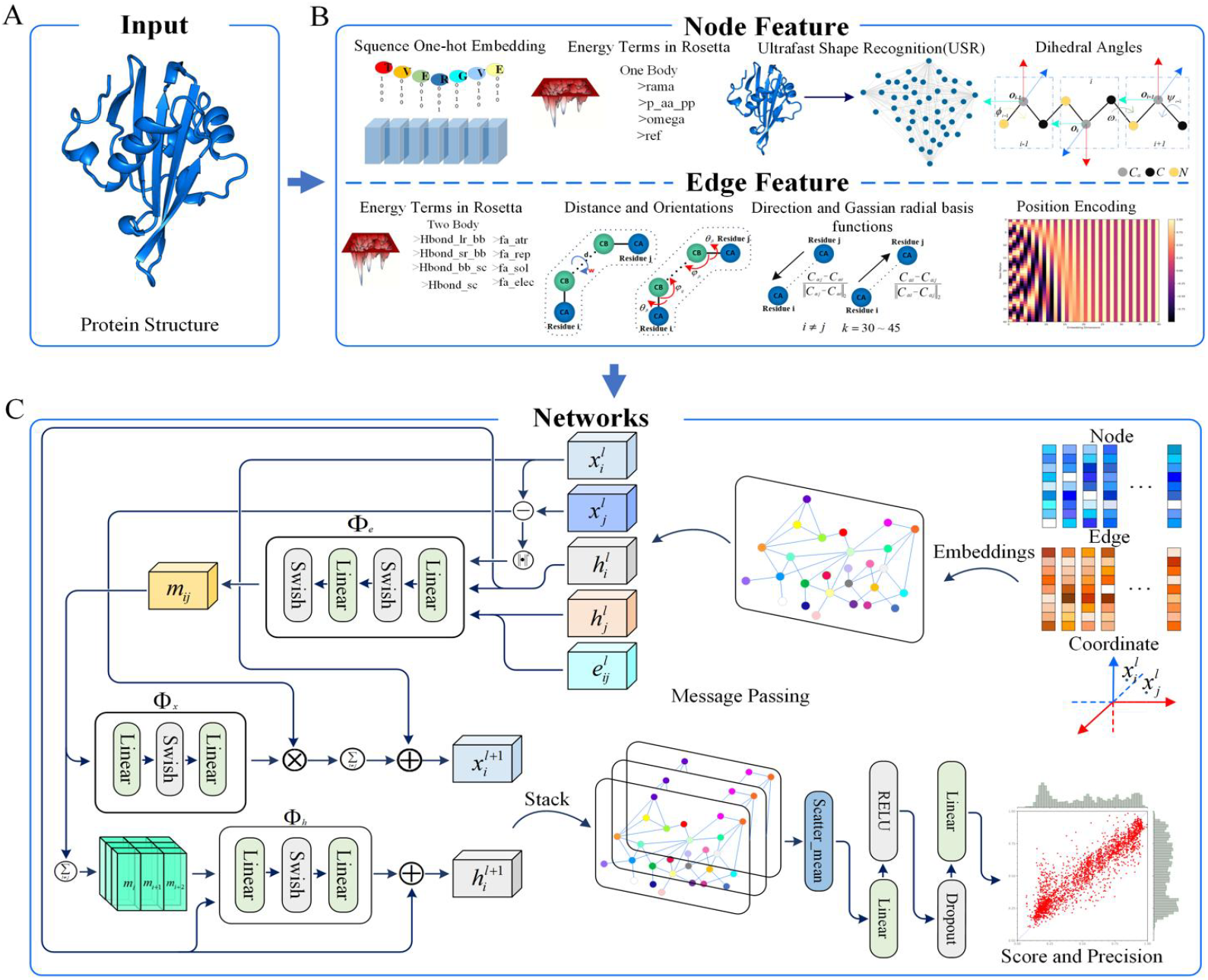
Pipeline of GraphGPSM. (A) Input protein structure. (B) Node and edge features representing the protein structure. Node features include one-hot encoding of residues, Rosetta one-body energy terms, residue-level USR, and dihedral angles. Edge features include Rosetta two-body energy terms, distance and orientations between residues, Gaussian radial basis functions encoding distance and direction, and residue sinusoidal position encoding. (C) Architecture of the graph neural network. The entire network embeds the feature information encoded in part B into the nodes and edges of an undirected graph and uses the designed message passing mechanism to update and transfer messages between layers. A multilayer perceptron is connected after the last layer to read the final score.

### 2.1 Data preparation

The protein dataset of GraphGPSM is constructed from PDB and AlphaFold Protein Structure Database [15, 16]. We screen 15,054 structures from the PDB (March 19, 2022) through the PISCES server [17], in which the selection criteria are sequence similarity less than 35%, structure length between 50 and 400, and minimum resolution of 2.5 Å. In addition, we select 1,118 human proteins from the AlphaFold Protein Structure Database (released in March 2022). The selection criteria are plDDT>90, not resolved in the PDB, sequence similarity less than 35%, and sequence similarity with the screened structure from the PDB database less than 35%. Finally, a total of 16,172 proteins are obtained.

For training GraphGPSM, three approaches are employed to generate about 150 structure models for each protein in the protein dataset: native perturbation, template modeling, and deep learning modeling. For native perturbation, we fine-tune the dihedral angle of the structure and fast relax the structure. Then, about 60 structure models are selected in accordance with the Rosetta energy of the perturbed structures. For template modeling, about 40 structure models are generated by RosettaCM [18] and Modeller [19] using different template structures and fragment libraries. For deep learning modeling, about 50 structure models are generated using our in-house method RocketX [20] by applying short-, medium-, and long-range constraints with different constraint weights. Lastly, the decoy structures generated by all methods are clustered, and similar structures are filtered out, resulting in about 1,473,706 structure models. All structure models are used to train the neural network.

### 2.2 Feature design

As shown in Figure 1(B), the protein structure model is represented by a graph, which contains node and edge features. Node features (i.e., one-hot encoding of sequences, Rosetta one-body energy terms, and residue-level USR) are used to represent information about each residue. Edge features (i.e., distance and orientations used in trRosetta, Rosetta two-body energy terms, and residue sinusoidal position encoding) are used to represent information about interactions between residues.

#### 2.2.1 Node features

To represent protein structures at the residue level, we employ four node features. First, each residue of the structure is represented as a 20D tensor through one-hot encoding to record the sequence information of the structure. Then, to describe the physicochemical effects of individual residues, we normalize the one-body terms of the centroid energy term in Rosetta to node features. One-body terms include rama, p_aa_pp, omega, and ref [7]. The residue-level USR designed in our previously developed DeepUMQA [11] can characterize the topological relationship between residues and the overall structure. Here, we utilize the USR feature to characterize protein structure information [21]. Briefly, for the current residue *r_i_*, the residue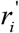 farthest from *r_i_* and the residue 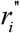 that farthest from 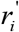 are first selected, and then the first moments of distances from all residues to the three residues are calculated as residue-level topological feature. Lastly, we embed the dihedral angles *ϕ_i_*, *φ_i_*, *ω_i_*, formed by adjacent planes to describe the relationship of adjacent residues. We use the sine and cosine of these three dihedral angles as node feature.

#### 2.2.2 Edge features

The edge features in the graph describe the relationship between different residues. Here, we employ four edge features. First, the Rosetta two-body energy terms are introduced to describe the physicochemical interaction between two residues, including fa_atr, fa_rep, fa_sol, fa_elec, hbond_lr_bb, hbond_sr_bb, hbond_bb_sc, and hbond_sc [7]. Then, inter-residue distance and orientations are used to describe the geometric relationship between residues. Gaussian radial basis function coding is an efficient machine learning coding method for mapping data into high-dimensional feature spaces. Thus, we use Gaussian radial basis functions to encode the distance of the main chain *C_α_* atoms and the direction of neighboring residues. Specifically, for each residue *i*, the nearest 30 residues are found in accordance with the distance between _*Cα*_ atoms. For each residue *j* in the 30 nearest residues, the unit vector *C_αi_* − *C_αj_* / *║C_αi_* − *C_αi_║*_2_ is embedded in the graph edges to represent the extension direction of the main chain. In this unit vector, *C_αi_* − *C_αi_* represents the direction from *C_αj_* to *C_αj_*, and *║C_αi_* − *C_αi_║*_2_ is the distance between _*Cα*_ atoms encoded by Gaussian radial basis functions. Lastly, we represent the positional relationship between these neighboring residues through sinusoidal positional encoding.

### 2.3 Network architecture

The network architecture of GraphGPSM is shown in Figure 1(C). It uses a message passing mechanism [22], in which nodes are updated with the information from neighboring nodes and edges during the propagation of each graph. On this basis, we have improved it and introduced the concept of EGNNs [23]. Traditionally, a graph *G* = (*V, E*) with nodes *υ_i_ ϵ V* and edges *e_ij_* ϵ *E*. Here, in addition to the feature node embeddings *h_i_* ϵ *R^nf^*, we also consider an n-dimensional coordinate *x_i_* ϵ *R ^nf^* associated with each of the graph nodes. Our model preserves equivariance to rotations and translations on this set of coordinates *x_i_* and equivariance to permutations on the set of nodes *V* in the same fashion as graph neural networks (GNNs), where *x_i_* denotes the *C_α_* atom coordinates of each residue. More specifically, the EGNN we use takes the set of node embeddings 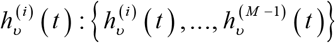, coordinate embedding *x_i_* (*t*) : {*x*_0_ (*t*), …, *x_M−1_* (*t*)}, and edge information *E* : { *e_ij_*} as input and outputs a transformation on 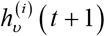 and *x_i_*(*t* +1). The equations that define this layer are the following:

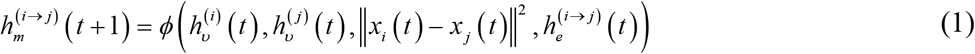

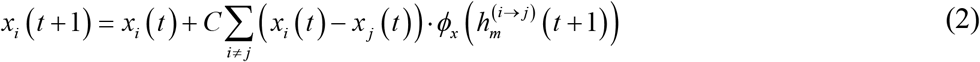

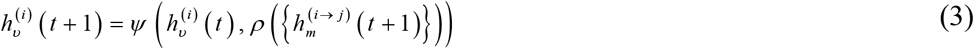

Where 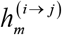 indicates that the message is passed from node *i* to node *j*; 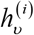 represents the embedding of node *i*; 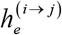 represents the feature embedding of edge (*i* → *j*) ; *ϕ* is the message function defined on each edge, which generates a message by combining the features on the edge with the features of the nodes at both ends. The relative squared distance between two coordinates *║x_i_(t)* − *x_j_ t)║*^2^ is input into the message *ϕ*. The aggregation function *ρ* aggregates the message received by the node. The update function *φ* combines the aggregated message and the features of the node itself to update the node features. In Equation 2, the position of each atom is updated by the weighted sum of all relative differences (*x_i_* − *x_j_)_⩝j_*. The weights of this sum are provided as the output of the function *ϕ_x_* : *R^nf^* → *R*^1^ that takes the edge embedding 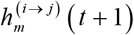 from the previous edge operation as input and outputs a scalar value. C is set to 1 / (*M* − 1), which divides the sum by its number of residues.

### 2.4 Network training

During model training, in each epoch, we divide the constructed data into training and validation sets in a ratio of 8:2. We use 50 graph propagation steps and regress on the scores of candidate structures. We implement the network by using PyG and PyTorch and train the neural network via the Adam optimizer with the following parameters: b1=0.9 and b2=0.999. We set the learning rate to 0.001 and the batch size to 8 and use mean squared error as the loss function. Training a model on our in-house dataset takes about a week on a Tesla V100 GPU.

## 3 Results

### 3.1 Global scoring performance on previous CASP and CAMEO targets

The performance of GraphGPSM is tested on the CASP13, CASP14, and CAMEO test sets and compared with that of the REF2015 energy function in the Rosetta modeling suite [7] and state-of-the-art model quality assessment methods, including ModFOLD7 [24], ProQ3D [25], VoroMQA-A [26], DeepAccNet [13], ModFOLD8 [27], QDeep [28], QMEAN [29], ProQ3 [30], and ModFOLD6 [31]. The CASP13 dataset contains 81 targets, each with approximately 150 structure models, for a total of 12,150 models. The CASP14 dataset contains 69 targets, each with approximately 149 structure models, for a total of 10,281 models. The CAMEO dataset (7 May to 30 July 2022) contains 181 targets, each with about 10 structure models, for a total of 1,810 models. The results of all lDDT-based scoring methods come from the CASP (https://predictioncenter.org/casp15/zscore_EMA.cgi) official website and the CAMEO (https://www.cameo3d.org/quality-estimate/) official website. The performance of GraphGPSM and the comparison methods on the CASP13, CASP14, and CAMEO test sets is shown in Figure 2, and the detailed results on each target are listed in Supplementary Tables S1, S2, and S3.

**Figure 2.**
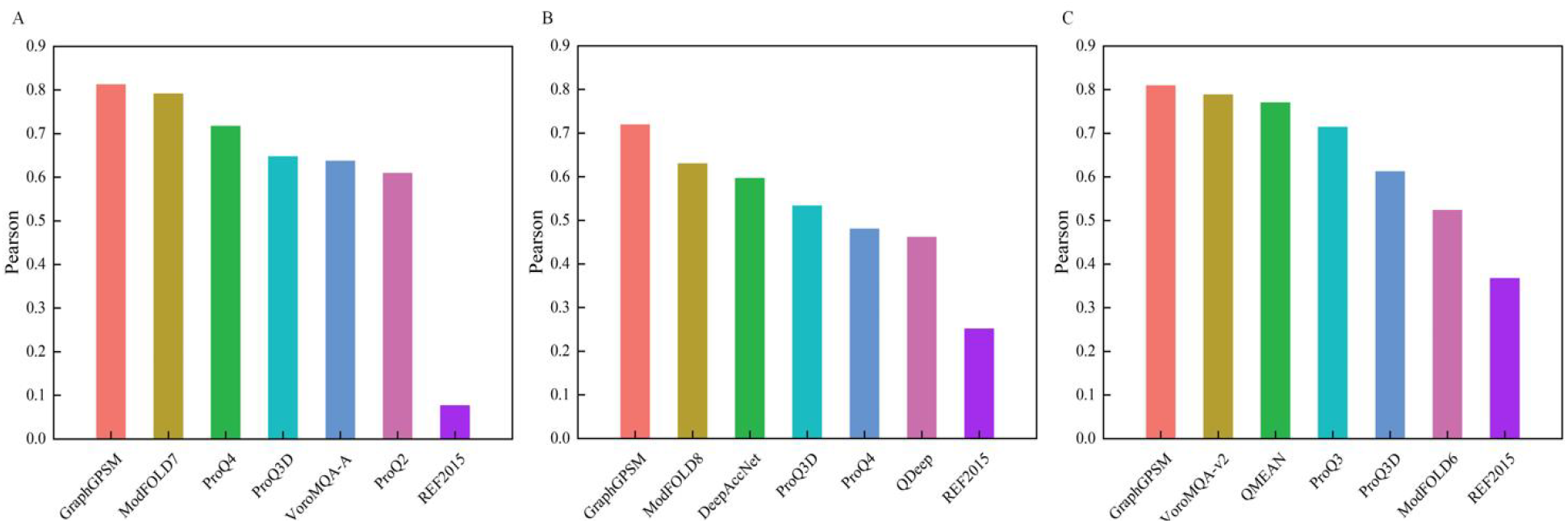
Average Pearson correlation coefficients of GraphGPSM and the comparison methods on (A) CASP13 (12,150 models), (B) CASP14 (10,281 models), and (C) CAMEO (1,810 models) test sets.

Here, we use the Pearson correlation between the global scoring of the protein structure model and the actual accuracy of the model (i.e., TM-score) as a measure of the performance of the method. GraphGPSM outperforms all the comparison methods on the CASP13, CASP14, and CAMEO test sets. On the CASP13 test set, the average Pearson correlation coefficient of GraphGPSM is 0.812, which is 2.4% higher than that of the second-ranked method, ModFold7. On the CASP14 test set, the average Pearson correlation coefficient of GraphGPSM is 0.719, which is 21% higher than that of the second-ranked method, ModFold8. On the 3-month CAMEO test set, the average Pearson correlation coefficient of GraphGPSM is 0.809, which is 2.7% higher than that of the second-ranked method, VoroMQA-v2. Compared with the scoring of the advanced lDDT-based model quality assessment methods, GraphGPSM’s scoring has a stronger correlation with the real quality score of the model. GraphGPSM also outperforms REF2015 significantly on all test sets. REF2015 is a scoring function used to calculate the interactions of all atoms in a protein to guide protein modeling. Owing to the lack of protein-specific knowledge, its performance is relatively poor. GraphGPSM can make up for this defect to a certain extent.

### 3.2 Blind test global scoring performance on CASP15

In this section, we show the performance of GraphGPSM on CASP15. As a plug-in algorithm, GraphGPSM is embedded into the participating server, GuijunLab-Threader (group number: 282), to participate in estimating model accuracy prediction [36]. The competition result shows that GraphGPSM has entered the top 10 global scoring servers. We compare the performance of all top-ranked servers on all 39 targets of CASP15, and the results are shown in Table 6. The detailed results on each target are presented in Supplementary Table S4. The data of all methods are obtained from the CASP official website (https://predictioncenter.org/download_area/CASP15/predictions/QA/). The average TM-score predicted by GraphGPSM is 0.730, which is closer to the real value of 0.715. The average Pearson correlation coefficient of the GraphGPSM scoring and the real TM-score is 0.561, which is higher than that of ModFOLDdock but lower than that of MULTICOM_qa. Among the 39 targets scored by GraphGPSM, 14 targets have a Pearson correlation coefficient greater than 0.9. The average bias of GraphGPSM is 0.126, which is the lowest among all the top-ranked servers. Minimal mean deviations are achieved on 16 targets.

The blind test results suggest that although the average Pearson coefficient between the TM-score predicted by our method and the real TM-score is slightly lower, the average TM-score it predicts is closer to the real value of average TM-score. We take H1134, T1170, and H1137 as examples to analyze the performance of GraphGPSM. The analysis for these three targets is shown in Figure 3. For the target H1134, the average real TM-score is 0.844, the average TM-score predicted by GraphGPSM is 0.836, and the average TM-score predicted by MULTICOM_qa is 0.460. Although, on the Pearson correlation coefficient, GraphGPSM is 0.964, and MULTICOM_qa is 0.982, the mean deviation between the GraphGPSM scoring and the true TM-score is smaller, only 0.04. From the scatter plot, the score predicted by GraphGPSM is closer to the real TM-score. For large-scale proteins, such as targets T1170o and H1137, the prediction performance of GraphGPSM is also better than that of the top-ranked server.

**Figure 3.**
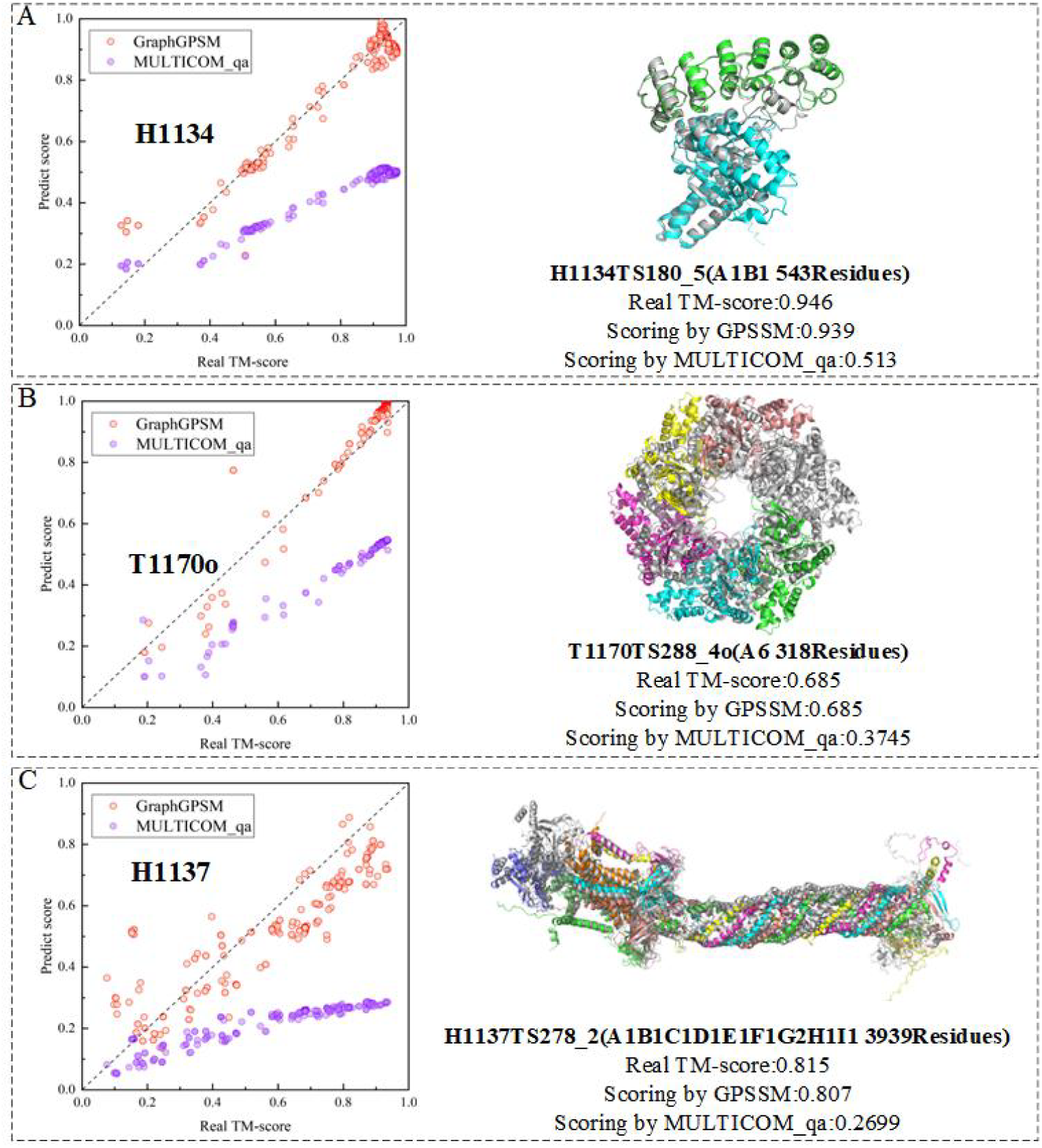
Case study for three targets of CASP15 (A) Target H1134. The average Pearson correlation coefficients of GraphGPSM and MULTICOM_qa are 0.836 and 0.460, respectively. The mean deviations of GraphGPSM and MULTICOM_qa are 0.040 and 0.386, respectively. (B) Target T1170o. The average Pearson correlation coefficients of GraphGPSM and MULTICOM_qa are 0.922 and 0.502, respectively. The mean deviations of GraphGPSM and MULTICOM_qa are 0.066 and 0.363, respectively. (C) Target H1137. The average Pearson correlation coefficients of GraphGPSM and MULTICOM_qa are 0.509 and 0.213, respectively. The mean deviations of GraphGPSM and MULTICOM_qa are 0.114 and 0.352, respectively.

### 3.3 Ablation studies

To analyze the impact of various features on the performance of GraphGPSM, we train 6 versions of GraphGPSM with the same dataset and the network architecture for different combinations of features. The performance of these 6 different versions of GraphGPSM is tested on the CASP13, CASP14, and CAMEO test sets. The sequence one-hot embedding and dihedral angle in node features and the positional encoding in edge features are used as the baseline version (version 1), and different features are combined on this basis to form different versions. Version 6 is a full version of GraphGPSM. The performance of different versions of GraphGPSM is shown in Figure 4, and the detailed results are listed in Supplementary Table S5.

**Figure 4.**
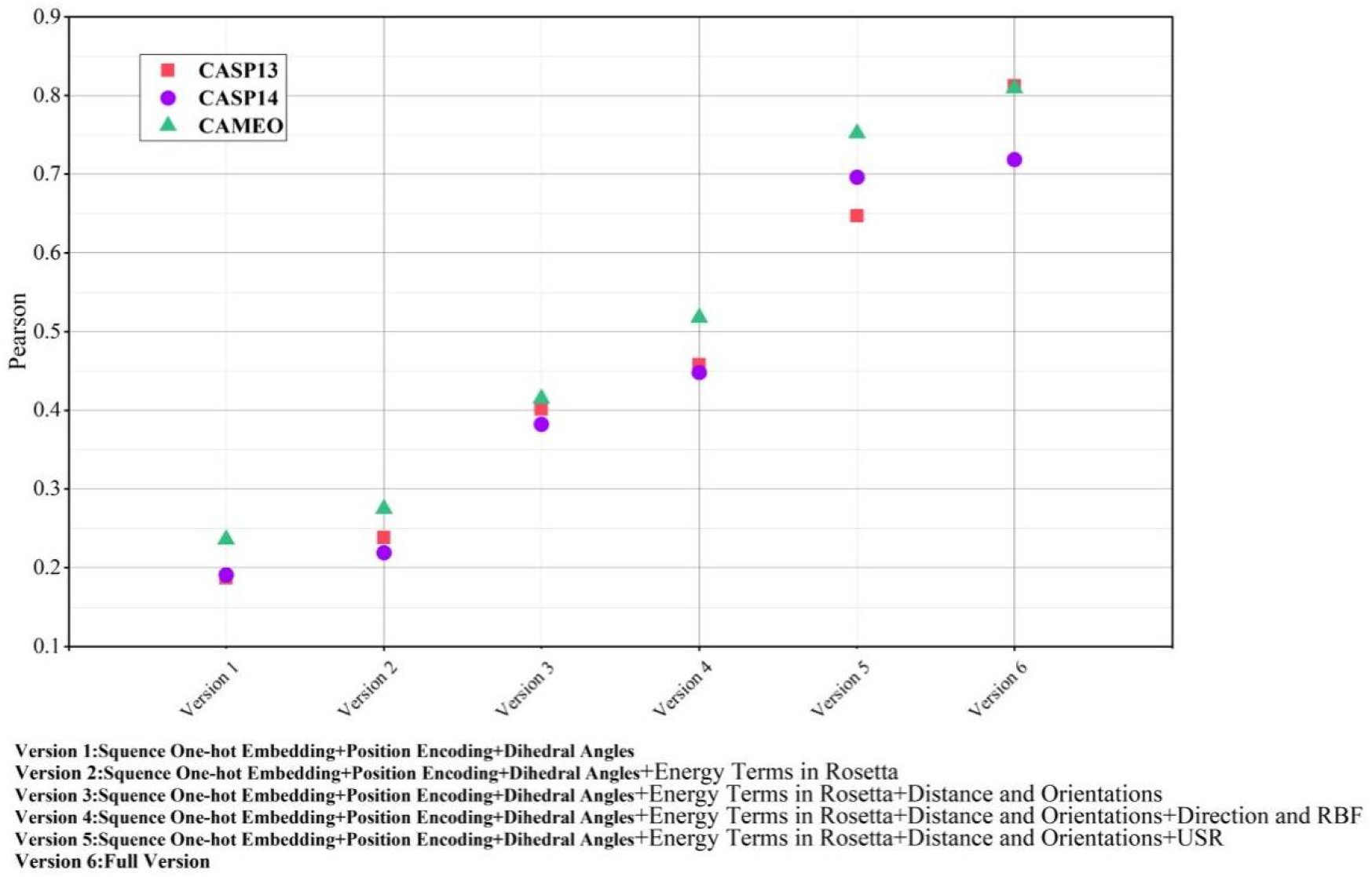
Performance of different versions of GraphGPSM on CASP13, CASP14, and CAMEO test sets.

The average Pearson correlation coefficients of version 1 on the three test sets are 0.187, 0.191, and 0.236, and those of version 2 (adding Rosetta energy terms on the basis of version 1) are increased to 0.238, 0.219, and 0.275. We add the inter-residue distance and orientations to form version 3 on the basis of version 2. The average Pearson correlation coefficients of version 3 on the three test sets are 0.401, 0.382, and 0.415; the performance is improved significantly. This result suggests that inter-residue geometries provide important structural information. Versions 4 and 5 add the distance and direction encoded by Gaussian radial basis functions and the residue-level USR features, respectively, on the basis of version 3. The performance of both versions is improved significantly. Especially, the performance of version 5 on the three test sets is improved by 61.3%, 82.1%, and 81.2%. Hence, the distance and direction encoded by Gaussian radial basis functions and residue-level USR contribute significantly to the performance of GraphGPSM. Version 6 adds the distance and direction encoded by Gaussian radial basis functions and residue-level USR on the basis of version 3, which is a significant improvement in comparison with versions 4 and 5. This suggests that the distance and direction encoded by Gaussian radial basis functions and residue-level USR can play complementary roles. In general, features based on geometric information (e.g., residue-level USR and distance and direction encoded by Gaussian radial basis functions) improve the performance more significantly than Rosetta energy terms do.

### 3.4 GraphGPSM-guided protein structure modeling

In this section, we further verify the performance of GraphGPSM. We integrate GraphGPSM into the Rosetta ClassicAbinitio protocol [32] and combine it with the Rosetta scoring function REF2015 [7] to guide protein structure modeling. The experimental results show that GraphGPSM improves the modeling performance of regular proteins by 67% compared with the Rosetta scoring function REF2015. In terms of the performance in modeling orphan and multi-domain proteins, the use of the structure predicted by AlphaFold2 as the initial model leads to 13.1% and 7.2% improvement, respectively. This result shows that simply adding GraphGPSM scoring to a classical scoring function can improve the modeling performance and that GraphGPSM scoring is effective. The above modeling is achieved by fragment assembly [33]. Then, the conformation is scored using GraphGPSM and the Rosetta energy function REF2015 [7] and decided to be accepted or rejected in accordance with the Metropolis criterion [34]. The final model is generated by iterating the above process.

#### Validation of the ability to guide protein modeling

We use the above algorithm to predict models for 484 proteins of our in-house IPTDFold [10] test set. To analyze the effect of GraphGPSM in protein modeling, we design a comparative experiment that removes the GraphGPSM scoring model from the above algorithm, that is, only REF2015 [7] is used to guide protein modeling. The comparison of both methods is shown in Figure 5(A), and the detailed results of each protein are listed in Supplementary Table S6. After the GraphGPSM scoring model is added, the accuracy of most proteins is improved significantly, and the average TM-score is increased by 66.5%. Among all tested proteins, 275 proteins show an increase in TM-score of more than 0.1, and 41 proteins show an increase of more than 0.2 after the addition of GraphGPSM. Here, we simply verify whether GraphGPSM can help guide protein structure modeling and therefore do not introduce additional constraints (e.g., distance and orientations) to design an elegant folding method. The experimental results show that GraphGPSM can be further used in the study of protein folding, which is faster and more accurate, while most of the existing end-to-end modeling methods can only obtain the final static structure.

**Figure 5.**
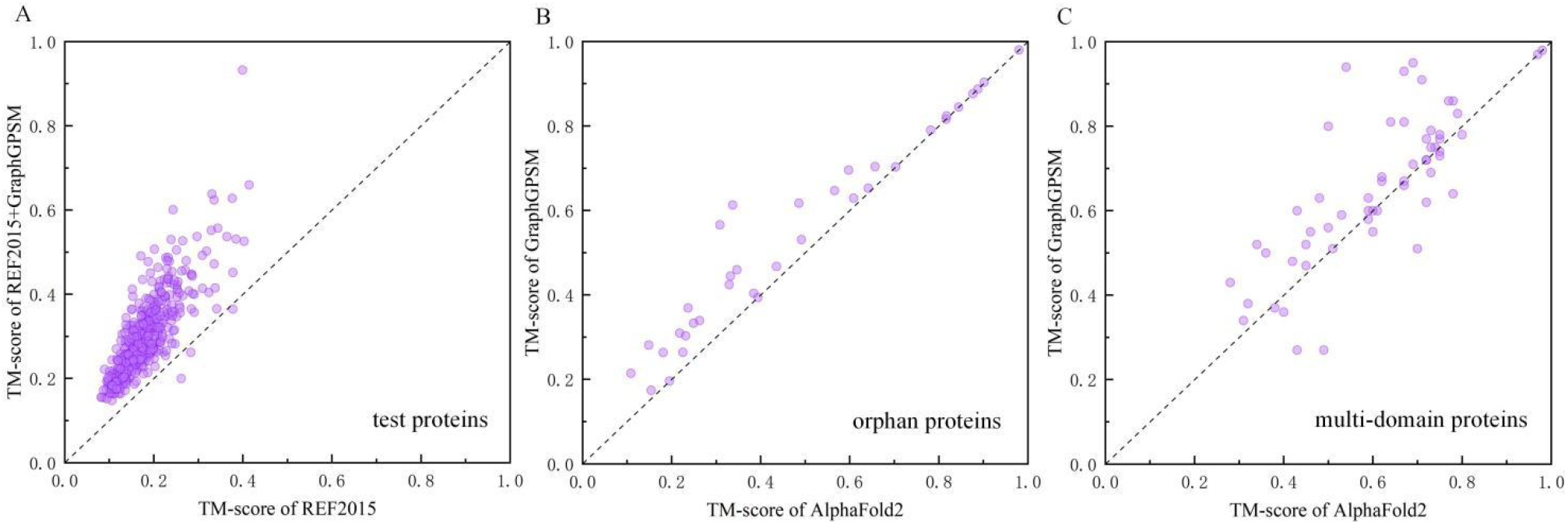
(A) Comparison of modeling with REF2015 energy function and modeling with both REF2015 energy function and GraphGPSM scoring model on 484 test proteins. (B) Comparison of TM-score of the initial model predicted by AlphaFold2 and the model improved by GraphGPSM on orphan proteins. (C) Comparison of TM-score of the initial model predicted by AlphaFold2 and the model improved by GraphGPSM on multi-domain proteins.

#### The ability to model orphan proteins

We randomly select 35 orphan proteins from the AlphaFold Protein Structure Database [16] to test the ability of GraphGPSM to model orphan proteins. For these orphan proteins, the model predicted by AlphaFold2 is used as the initial structure of the above GraphGPSM modeling algorithm to repredict the structure models. The results of AlphaFold2 and GraphGPSM are summarized in Table 1, and the detailed results of each protein are listed in Supplementary Table S7. Figure 5(B) visually reflects the comparison between AlphaFold2 and GraphGPSM. The average TM-score of the initial structure models predicted by AlphaFold2 is 0.478, and that of the structures improved by GraphGPSM is increased by 13.1%. AlphaFold2 correctly folds (TM-score ≥0.5) 14 of the 35 orphan proteins, which increase to 18 after the improvement with GraphGPSM. The TM-score of the proteins improved by GraphGPSM increases by more than 0.1 on 8 proteins and by more than 0.2 on 2 proteins. For the proteins with TM-score less than 0.7, the average TM-score increases by 23.9% after improvement; for the proteins with TM-score greater than 0.7, the average TM-score only increases by 0.1%. From Figure 5(A), a TM-score of 0.7 is a watershed. We can conclude that although the end-to-end modeling method has achieved very good results, if coevolutionary information is lacking, its modeling accuracy is still not enough. We may need a better scoring model to guide the protein structure modeling or quality ranking, which also shows that the scoring model and the end-to-end modeling method have a complementary improvement effect.

**Table 1.** Performance comparison with top-ranked servers on all targets of CASP15.

| Method | Average TM-score | Average Pearson | Average Bias |
| --- | --- | --- | --- |
| GraphGPSM | **0.730** | 0.633 | **0.126** |
| MULTICOM_qa | 0.485 | **0.715** | 0.258 |
| ModFOLDdock | 0.515 | 0.636 | 0.241 |
| ModFOLDdockR | 0.666 | 0.635 | 0.165 |
| Venclovas | 0.449 | 0.494 | 0.339 |
| Manifold | 0.582 | 0.541 | 0.179 |
| Bhattacharya | 0.387 | 0.474 | 0.361 |
| *Real value | 0.716 | — | — |

\*Real value represents the real average TM-score of all targets in CASP15.
The data of all methods are obtained from the CASP official website (https://predictioncenter.org/).

#### The ability to model multi-domain proteins

We randomly select 57 multi-domain proteins from the SADA [35] dataset to test the ability of GraphGPSM to model multi-domain proteins. For these multi-domain proteins, the model predicted by AlphaFold2 is used as the initial structure of the above GraphGPSM modeling algorithm to repredict the structure models. The results of AlphaFold2 and GraphGPSM are summarized in Table 2, and the detailed results of each protein are listed in Supplementary Table S8. Figure 5(C) visually reflects the comparison between AlphaFold2 and GraphGPSM. The average TM-score of the structures predicted by AlphaFold2 is 0.611, while the average TM-score of the improved structures is increased by 7%. 41 proteins are correctly folded by AlphaFold2, and 3 proteins exhibit a TM-score greater than 0.8. After GraphGPSM improvement, 47 proteins are correctly folded, and 12 proteins have a TM-score greater than 0.8. The TM-score of the proteins improved by GraphGPSM increases by more than 0.1 on 12 proteins and by more than 0.2 on 5 proteins. For the proteins with TM-score less than 0.5, the average TM-score increases by 15% after improvement; for the proteins with TM-score greater than 0.5, the average TM-score increases by 5.5%. Figure 6 shows the structural changes of four multi-domain proteins before and after improvement. The model accuracy is improved significantly after GraphGPSM improve. The reason is that the single-domain accuracy of the AlphaFold2 prediction model is high, but the interdomain orientation is inaccurate. On the contrary, GraphGPSM improvement can effectively adjust the interdomain orientation to increase the overall accuracy of the model.

**Table 2.** Results of AlphaFold2 and GraphGPSM on orphan proteins.

| Method | TM-score | #TM$\geq 0.5$ | #TM$\geq 0.6$ | #TM$\geq 0.7$ | #TM$\geq 0.8$ | P-value |
| --- | --- | --- | --- | --- | --- | --- |
| AlphaFold2 | 0.478 | 14 | 12 | 9 | 7 | NA |
| GraphGPSM | 0.541 | 18 | 16 | 10 | 7 | 2.752E-06 |

#TM $\geq 0.5$ , $\geq 0.6$ , $\geq 0.7$ , $\geq 0.8$ are the numbers of the predicted models with TM-score $\geq 0.5$ , $\geq 0.6$ , $\geq 0.7$ , $\geq 0.8$ , respectively.
P-value presents the results of the Wilcoxon signed-rank test.

**Figure 6.**
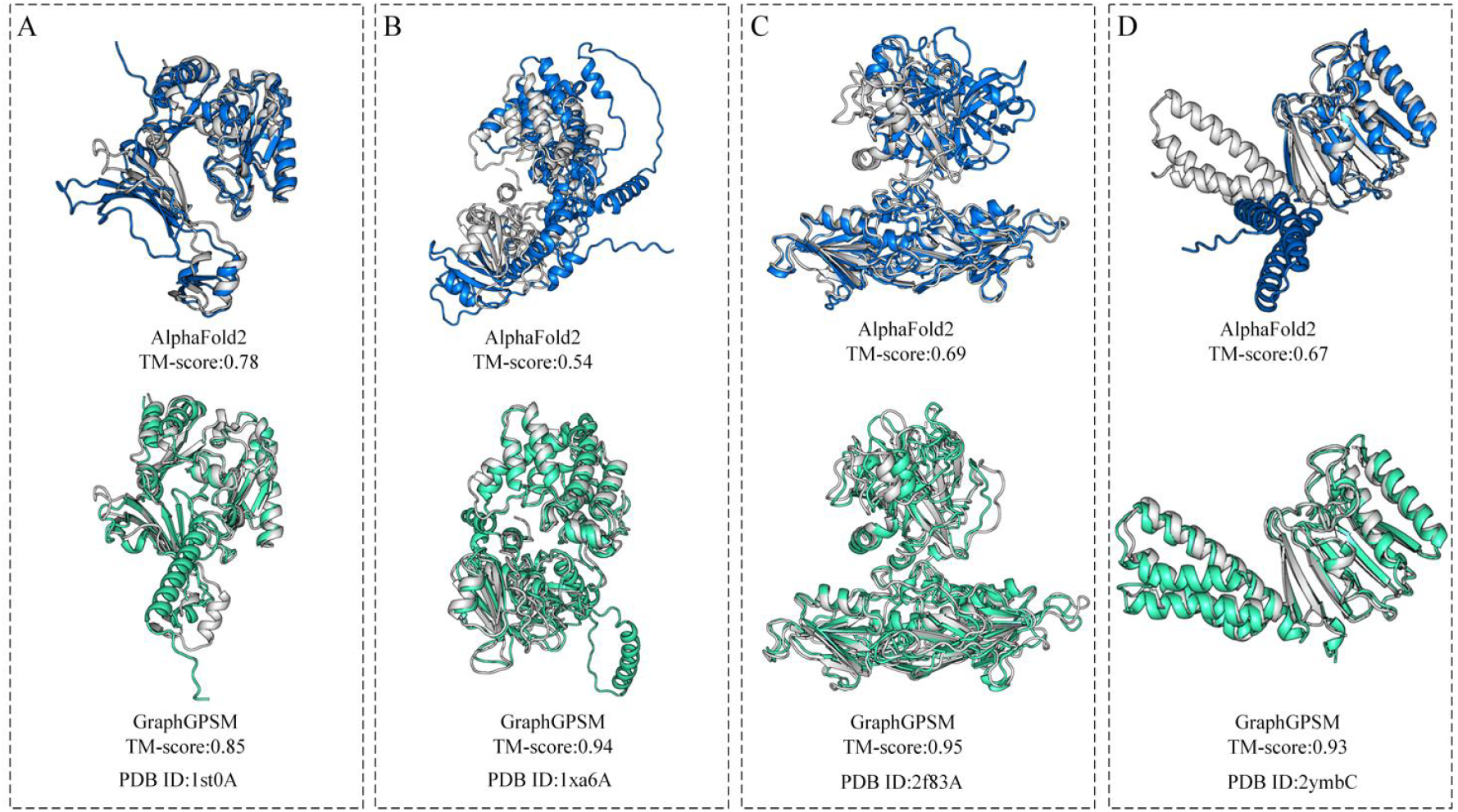
Four examples of multi-domain protein improvement by GraphGPSM. Gray is the native structure, blue is the initial structure model predicted by AlphaFold2, and green is the structure model improved by GraphGPSM.

**Table 3.** Results of AlphaFold2 and GraphGPSM on multi-domain proteins.

| Method | TM-score | #TM≥0.5 | #TM≥0.6 | #TM≥0.7 | #TM≥0.8 | P-value |
| --- | --- | --- | --- | --- | --- | --- |
| AlphaFold2 | 0.611 | 41 | 33 | 21 | 3 | NA |
| GraphGPSM | 0.656 | 47 | 37 | 24 | 12 | 0.002 |

#TM ≥ 0.5, ≥ 0.6, ≥ 0.7, ≥ 0.8 are the numbers of the predicted models with TM-score ≥ 0.5, ≥ 0.6, ≥ 0.7, ≥ 0.8, respectively. P-value presents the results of the Wilcoxon signed-rank test.

**Table 4.** Comparison of the model selected by GraphGPSM and the first model of AlphaFold2.

| Method | Average TM-score | #Best model | #Better than other |
| --- | --- | --- | --- |
| AlphaFold2 | 0.859 | 21/66 | 16/66 |
| GraphGPSM | 0.875 | 28/66 | 34/66 |

### 3.5 Model ranking ability

In this section, we further explore whether GraphGPSM can choose the best models for state-of-the-art protein prediction methods. We collect the top-5 models for 66 targets predicted by AlphaFold2 from the official website of CASP14. We score these structures by using GraphGPSM and select the top-scoring models to compare with the top-ranked models selected by AlphaFold2 self-assessment. The comparison results are summarized in Table 5 and Figure 7, and the detailed results for each target are shown in Supplementary Table S9. For the 66 targets of AlphaFold2, the average TM-score of the model selected by GraphGPSM is 0.875, which is higher than that (0.859) of the first model of AlphaFold2. GraphGPSM selects the best model on 28 of the 66 targets, and those of AlphaFold2 is 21. The model selected by GraphGPSM is more accurate than the first model of AlphaFold2 on 34 targets, and the first model of AlphaFold2 is more accurate on 16 targets. Figure 7 also demonstrates that the GraphGPSM-selected models with mean and median TM-score are closer to the real best model. Therefore, GraphGPSM can select a more accurate model from the models predicted by AlphaFold2.

**Figure 7.**
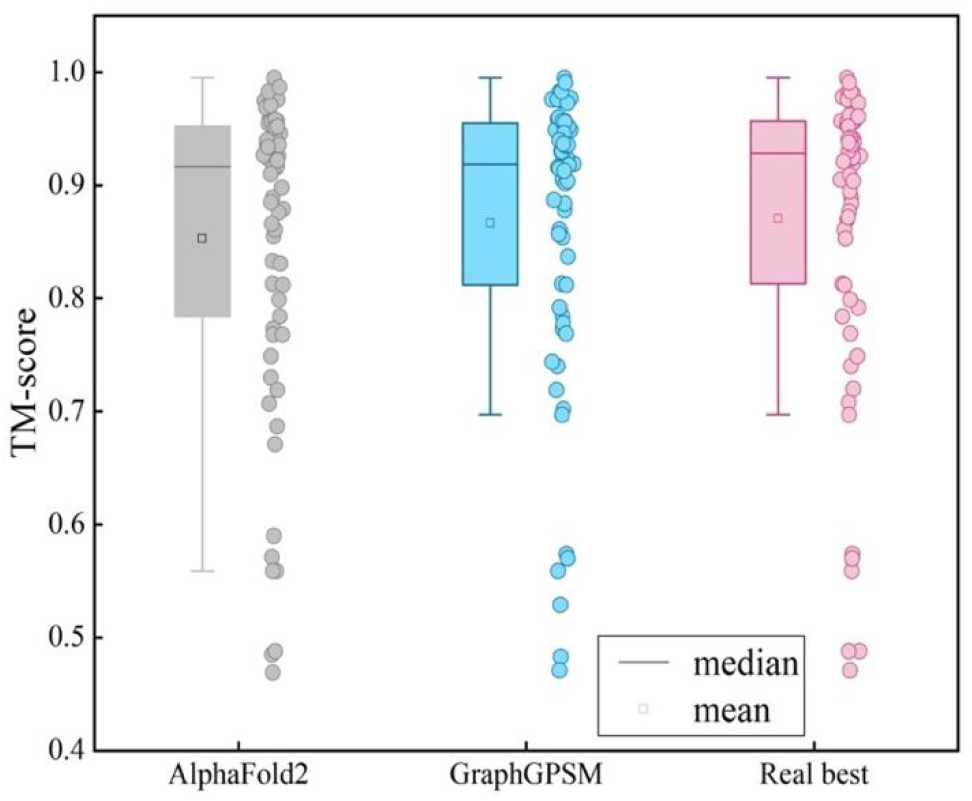
Boxplots and distributions of AlphaFold2’s first model, the best model selected by GraphGPSM, and the real best model on 66 CASP14 targets.

## 4 Conclusion

We develop a protein structure scoring model based on GNN, called GraphGPSM. Residue-level USR, Gaussian radial basis function-encoded distance and orientations, dihedral angle, one-hot encoding of residues, sinusoidal position encoding of amino acid sequence, Rosetta energy term, distance, and orientations are used to characterize the model information of proteins. The information of these features is fed into the GNN to predict the global score of the protein structure model. The experimental results show that the performance on the CASP13, CASP14, and CAMEO test sets is better than that of the state-of-the-art single-model quality methods, including ModFOLD7, ProQ3D, VoroMQA-A, DeepAccNet, ModFOLD8, QDeep, QMEAN, ProQ3, ProQ3D, and ModFOLD6. For orphan and multidomain proteins, GraphGPSM can further refine the structure predicted by AlphaFold2, and adding GraphGPSM on the basis of the Rosetta energy function can help guide protein structure modeling. Moreover, GraphGPSM can select more accurate structure models predicted by AlphaFold2. GraphGPSM’s performance on CASP15 also makes it into a top-ranked server. At present, GraphGPSM’s ability to capture the interface relationship of complex protein structures is still in continuous improvement, and there is still much room for improvement in the performance of scoring complexes. In the future, scoring the global quality of complexes will be a crucial task.

## Supporting information

Support Information

## Key points

- We designed a protein structure scoring model (GraphGPSM) based on graph neural networks. In GraphGPSM, geometric features and physicochemical energy terms are used together to characterize protein structures; and a well-designed message passing mechanism enables good information updating and passing on nodes and edges.
- The experimental results show that, compared with the unified field energy model and the IDDT-based scoring model, the GraphGPSM score has a stronger correlation with the real global score of the protein structure, and it can help guide protein structure modeling. The accuracy of the model picked by GraphGPSM is better than AlphaFold2’s self-evaluation.
- For orphan proteins and multi-domain proteins, GraphGPSM can further refine the structures predicted by AlphaFold2. In CASP15, GraphGPSM entered the top-ranked server for EMA prediction.

## Supplementary Data

Supplementary data are available online at BIB.

## Data and code availability

All data needed to evaluate the conclusions are present in the paper and the Supplementary Materials. The web server of GraphGPSM is freely available at http://zhanglab-bioinf.com/GraphGPSM/.

