## Supplementary material for "GraphGPSM: a global scoring model for protein structure using graph neural networks": Support Information

### Supplementary Information

Table S1. Detailed information of the 81 target proteins in the CASP13 datasets. #Model represents the number of models.

| Target | Length | #Model | Pearson | Target | Length | #Model | Pearson |
|---|---|---|---|---|---|---|---|
| T0949 | 183 | 150 | 0.850 | T0987 | 405 | 150 | 0.857 |
| T0950 | 353 | 150 | 0.936 | T0988 | 204 | 150 | 0.820 |
| T0951 | 276 | 149 | 0.946 | T0989 | 246 | 150 | 0.741 |
| T0953s1 | 72 | 150 | 0.820 | T0990 | 552 | 150 | 0.550 |
| T0954 | 350 | 150 | 0.934 | T0991 | 118 | 150 | 0.454 |
| T0955 | 41 | 150 | 0.886 | T0992 | 126 | 150 | 0.896 |
| T0957s1 | 163 | 150 | 0.824 | T0993s1 | 269 | 150 | 0.546 |
| T0957s2 | 164 | 150 | 0.932 | T0993s2 | 109 | 150 | 0.614 |
| T0958 | 96 | 150 | 0.897 | T0994 | 585 | 150 | 0.875 |
| T0959 | 189 | 150 | 0.840 | T0995 | 330 | 150 | 0.753 |
| T0960 | 384 | 150 | 0.909 | T0996 | 848 | 150 | 0.801 |
| T0961 | 505 | 150 | 0.908 | T0997 | 228 | 150 | 0.874 |
| T0962 | 220 | 150 | 0.898 | T0998 | 166 | 150 | 0.694 |
| T0963 | 372 | 150 | 0.799 | T0999 | 1589 | 143 | 0.838 |
| T0964 | 184 | 150 | 0.928 | T1000 | 523 | 150 | 0.951 |
| T0965 | 334 | 150 | 0.724 | T1001 | 140 | 150 | 0.858 |
| T0966 | 494 | 150 | 0.914 | T1002 | 270 | 150 | 0.858 |
| T0967 | 81 | 150 | 0.845 | T1003 | 474 | 150 | 0.743 |
| T0968s1 | 126 | 150 | 0.822 | T1004 | 458 | 150 | 0.509 |
| T0968s2 | 116 | 150 | 0.925 | T1005 | 364 | 150 | 0.925 |
| T0969 | 487 | 150 | 0.732 | T1006 | 79 | 150 | 0.807 |
| T0970 | 97 | 150 | 0.822 | T1008 | 80 | 150 | 0.616 |
| T0971 | 186 | 150 | 0.871 | T1009 | 718 | 150 | 0.892 |
| T0973 | 146 | 150 | 0.97 | T1010 | 210 | 150 | 0.872 |
| T0974s1 | 72 | 150 | 0.821 | T1011 | 534 | 150 | 0.914 |
| T0974s2 | 95 | 150 | 0.927 | T1013 | 537 | 150 | 0.956 |
| T0975 | 343 | 150 | 0.802 | T1014 | 276 | 150 | 0.803 |
| T0976 | 252 | 150 | 0.817 | T1015s1 | 89 | 150 | 0.776 |
| T0977 | 566 | 150 | 0.761 | T1015s2 | 129 | 150 | 0.860 |
| T0978 | 416 | 150 | 0.796 | T1016 | 203 | 150 | 0.582 |
| T0979 | 98 | 150 | 0.782 | T1017s1 | 111 | 150 | 0.786 |
| T0980s1 | 111 | 150 | 0.857 | T1017s2 | 129 | 150 | 0.704 |
| T0980s2 | 52 | 150 | 0.766 | T1018 | 334 | 150 | 0.931 |
| T0981 | 640 | 150 | 0.834 | T1019s1 | 58 | 150 | 0.656 |
| T0982 | 283 | 150 | -0.002 | T1019s2 | 88 | 150 | 0.912 |
| T0983 | 245 | 150 | 0.901 | T1020 | 577 | 150 | 0.99 |
| T0984 | 752 | 150 | 0.975 | T1021s1 | 149 | 150 | 0.959 |
| T0985 | 863 | 150 | 0.8 | T1021s2 | 354 | 150 | 0.932 |
| T0986s1 | 96 | 150 | 0.801 | T1021s3 | 295 | 150 | 0.716 |
| T0986s2 | 155 | 150 | 0.717 | T1022s1 | 229 | 150 | 0.722 |
| T1022s2 | 529 | 150 | 0.892 | | | | |

Table S2. Detailed information of the 69 target proteins in the CASP14 datasets. #Model represents the number of models.

| Target | Length | #Model | Pearson | Target | Length | #Model | Pearson |
|---|---|---|---|---|---|---|---|
| T1024 | 408 | 150 | 0.782 | T1062 | 35 | 150 | 0.316 |
| T1025 | 268 | 150 | 0.182 | T1064 | 106 | 150 | 0.529 |
| T1026 | 172 | 150 | 0.784 | T1065s1 | 127 | 150 | 0.745 |
| T1027 | 168 | 150 | 0.713 | T1065s2 | 98 | 150 | 0.707 |
| T1028 | 316 | 150 | 0.483 | T1067 | 264 | 150 | 0.487 |
| T1029 | 125 | 150 | 0.044 | T1070 | 335 | 150 | 0.819 |
| T1030 | 273 | 150 | 0.713 | T1072s1 | 101 | 150 | 0.966 |
| T1031 | 95 | 150 | 0.912 | T1073 | 255 | 150 | 0.963 |
| T1032 | 284 | 150 | 0.695 | T1074 | 202 | 150 | 0.834 |
| T1033 | 100 | 150 | 0.903 | T1076 | 552 | 150 | 0.266 |
| T1034 | 156 | 150 | 0.400 | T1078 | 138 | 150 | 0.812 |
| T1035 | 102 | 150 | 0.974 | T1079 | 505 | 150 | 0.792 |
| T1036s1 | 622 | 150 | 0.649 | T1080 | 922 | 144 | 0.934 |
| T1037 | 404 | 150 | 0.968 | T1082 | 97 | 150 | 0.818 |
| T1038 | 199 | 150 | 0.870 | T1083 | 98 | 150 | 0.306 |
| T1039 | 161 | 150 | 0.900 | T1084 | 73 | 150 | 0.103 |
| T1040 | 130 | 150 | 0.861 | T1087 | 93 | 150 | 0.921 |
| T1041 | 242 | 150 | 0.965 | T1088 | 226 | 150 | 0.840 |
| T1042 | 289 | 150 | 0.923 | T1089 | 404 | 150 | 0.421 |
| T1043 | 148 | 150 | 0.921 | T1090 | 193 | 150 | 0.802 |
| T1045s1 | 157 | 150 | 0.286 | T1091 | 863 | 150 | 0.924 |
| T1045s2 | 173 | 150 | 0.730 | T1092 | 426 | 150 | 0.930 |
| T1046s1 | 74 | 150 | 0.569 | T1093 | 631 | 150 | 0.920 |
| T1046s2 | 142 | 150 | 0.750 | T1094 | 496 | 150 | 0.711 |
| T1047s1 | 232 | 150 | 0.778 | T1095 | 665 | 150 | 0.934 |
| T1047s2 | 365 | 150 | 0.878 | T1096 | 464 | 150 | 0.702 |
| T1048 | 109 | 150 | 0.090 | T1098 | 538 | 150 | 0.986 |
| T1049 | 141 | 150 | 0.818 | T1099 | 262 | 150 | 0.834 |
| T1050 | 779 | 150 | 0.943 | T1100 | 338 | 150 | 0.803 |
| T1052 | 832 | 150 | 0.870 | T1101 | 318 | 150 | 0.308 |
| T1053 | 580 | 150 | 0.903 | T1061 | 949 | 150 | 0.687 |
| T1054 | 190 | 150 | 0.721 | | | | |
| T1055 | 148 | 150 | 0.818 | | | | |
| T1056 | 186 | 150 | 0.712 | | | | |
| T1057 | 287 | 150 | 0.521 | | | | |
| T1058 | 382 | 150 | 0.724 | | | | |
| T1060s2 | 298 | 150 | 0.921 | | | | |
| T1060s3 | 140 | 150 | 0.769 | | | | |

Table S3. Detailed information of the 198 target proteins in the CAMEO datasets. #Model represents the number of models.

| Target | Length | #Model | Pearson | Target | Length | #Model | Pearson |
|---|---|---|---|---|---|---|---|
| 7EQH_A | 228 | 14 | 0.855 | 8CU5_A | 179 | 7 | 0.748 |
| 7ER0_A | 307 | 9 | 0.795 | 8CUK_B | 363 | 7 | 0.985 |
| 7ERN_C | 273 | 9 | 0.868 | 8CTR_A | 296 | 9 | 0.606 |
| 7F0A_A | 280 | 9 | 0.810 | 7ZNX_A | 137 | 8 | 0.994 |
| 7F9H_A | 73 | 11 | 0.983 | 7WAW_A | 505 | 6 | 0.630 |
| 7MKU_A | 309 | 11 | 0.794 | 7W6B_A | 187 | 7 | 0.993 |
| 7OPB_D | 27 | 13 | 0.955 | 7W3R_A | 359 | 6 | 0.959 |
| 7POI_C | 295 | 10 | 0.964 | 7TZV_A | 188 | 7 | 0.888 |
| 7PRQ_B | 330 | 11 | 0.956 | 7TXC_E | 110 | 8 | 0.711 |
| 7PSG_C | 264 | 9 | 0.964 | 7TE3_A | 174 | 8 | 0.699 |
| 7SO5_H | 215 | 8 | 0.581 | 7R24_A | 159 | 8 | 0.391 |
| 7TZE_C | 232 | 9 | 0.959 | 7R20_B | 116 | 6 | 0.550 |
| 7TZG_D | 397 | 9 | 0.871 | 7QWT_A | 340 | 6 | 0.716 |
| 7X9E_A | 221 | 6 | 0.938 | 7QII_B | 45 | 12 | 0.720 |
| 7Z5P_A | 508 | 5 | 0.627 | 7OIO_A | 98 | 10 | 0.846 |
| 7ESH_A | 635 | 3 | 0.984 | 7OHZ_B | 277 | 10 | 0.220 |
| 7MQ4_A | 52 | 7 | 0.823 | 7FAU_B | 127 | 9 | 0.997 |
| 7MS2_A | 755 | 7 | 0.997 | 7EXX_A | 67 | 7 | 0.994 |
| 7R3P_A | 157 | 8 | 0.713 | 7EZB_A | 167 | 9 | 0.812 |
| 7R5Y_E | 381 | 6 | 0.979 | 7EZN_A | 177 | 9 | 0.971 |
| 7RKC_A | 68 | 7 | 0.999 | 7N45_A | 131 | 11 | 0.959 |
| 7SJL_A | 96 | 3 | 0.993 | 7OPT_A | 267 | 11 | 0.503 |
| 7SXB_A | 90 | 5 | 0.852 | 7PB9_A | 166 | 9 | 0.932 |
| 7W5M_A | 260 | 6 | 0.999 | 7PGF_D | 137 | 9 | 0.534 |
| 7WCJ_A | 443 | 4 | 0.953 | 7PGI_D | 138 | 9 | 0.864 |
| 7X4O_B | 131 | 7 | 0.997 | 7RE5_B | 117 | 7 | 0.723 |
| 7X8J_A | 361 | 5 | 0.992 | 7UW7_A | 76 | 12 | 0.975 |
| 7ZCL_B | 219 | 8 | 0.662 | 7VQW_A | 109 | 7 | 0.488 |
| 7ZGF_A | 407 | 7 | 0.989 | 7VRS_A | 441 | 14 | 0.860 |
| 7EUS_B | 285 | 10 | 0.947 | 7VTY_A | 89 | 10 | 0.905 |
| 7F5G_B | 116 | 9 | 0.656 | 7VU4_A | 108 | 9 | 0.691 |
| 7PCR_A | 551 | 10 | 0.540 | 7XH0_A | 259 | 10 | 0.855 |
| 7QRE_D | 103 | 10 | 0.838 | 8CSO_C | 230 | 10 | 0.415 |
| 7R1K_A | 179 | 8 | 0.823 | 8D1X_D | 468 | 11 | 0.877 |
| 7S69_A | 132 | 8 | 0.979 | 7XOI_A | 482 | 4 | 0.831 |
| 7TVC_B | 118 | 8 | 0.942 | 7XKG_A | 401 | 10 | 0.662 |
| 7TVY_B | 124 | 9 | 0.805 | 7XIF_D | 284 | 9 | 0525 |
| 7VGB_A | 681 | 5 | 0.958 | 7X0F_C | 521 | 7 | 0.971 |
| 7VRB_A | 49 | 10 | 0.756 | 7VMT_B | 140 | 8 | 0.990 |
| 7VRC_C | 169 | 10 | 0.977 | 7PCS_B | 243 | 7 | 0.625 |

| | | | | | | | |
|---|---|---|---|---|---|---|---|
| 7N8U_A | 262 | 9 | 0.579 | 7T7I_D | 237 | 10 | 0.986 |
| 7MO3_B | 41 | 12 | 0.513 | 7V67_A | 433 | 8 | 0.052 |
| 7MO1_B | 26 | 12 | 0.716 | 7VA8_A | 453 | 9 | 0.927 |
| 7MNY_D | 133 | 11 | 0.897 | 7X2E_A | 101 | 8 | 0.989 |
| 7MNV_B | 36 | 12 | 0.410 | 8CWU_B | 122 | 5 | 0.786 |
| 7MNK_A | 18 | 12 | 0.461 | 7XQV_B | 121 | 7 | 0.944 |
| 7MNI_C | 479 | 10 | 0.985 | 7U37_A | 46 | 5 | 0.940 |
| 7F4S_D | 227 | 9 | 0.990 | 7SAF_B | 57 | 8 | 0.952 |
| 7F4L_D | 194 | 9 | 0.403 | 7RY6_A | 435 | 8 | 0.937 |
| 7F15_B | 244 | 9 | 0.971 | 7PJJ_A | 824 | 4 | 0.927 |
| 7F08_N | 389 | 9 | 0.840 | 7P4L_B | 133 | 6 | 0.988 |
| 7B8B_A | 347 | 9 | 0.996 | 7P23_A | 230 | 7 | 0.780 |
| 7F5K_A | 191 | 10 | 0.980 | 7P22_A | 217 | 6 | 0.965 |
| 7OQ6_A | 406 | 8 | 0.615 | 7P20_A | 197 | 10 | 0.783 |
| 7OU3_A | 220 | 11 | 0.868 | 7OYL_A | 549 | 6 | 0.544 |
| 7OUU_B | 102 | 9 | 0.978 | 7OXL_A | 490 | 7 | 0.447 |
| 7QQA_A | 270 | 8 | 0.686 | 7FAV_A | 76 | 8 | 0.928 |
| 7QVB_B | 15 | 11 | 0.930 | 7BLF_A | 416 | 6 | 0.883 |
| 7R0R_A | 278 | 5 | 0.999 | 7BB8_A | 306 | 6 | 0.987 |
| 7U5O_C | 115 | 11 | 0.854 | 7ESG_A | 226 | 9 | 0.999 |
| 7V40_A | 432 | 7 | 0.934 | 7FCC_A | 141 | 10 | 0.924 |
| 7W7A_G | 228 | 6 | 0.833 | 7FDP_B | 397 | 8 | 0.975 |
| 7X77_A | 272 | 3 | 0.894 | 7Q1T_A | 22 | 12 | 0.989 |
| 7ZMN_K | 123 | 6 | 0.918 | 7R6R_A | 49 | 8 | 0.990 |
| 7ZMP_K | 123 | 8 | 0.682 | 7RFQ_A | 155 | 7 | 0.993 |
| 7ZMQ_K | 123 | 6 | 0.605 | 7RGW_A | 160 | 6 | 0.800 |
| 7ZMR_K | 123 | 8 | 0.736 | 7SL5_G | 235 | 8 | 0.984 |
| 7ZRO_A | 21 | 9 | 0.920 | 7SPY_A | 348 | 9 | 0.654 |
| 7DRI_A | 298 | 8 | 0.874 | 7T2S_A | 177 | 9 | 0.988 |
| 7PLN_A | 258 | 9 | 0.993 | 7UM4_A | 325 | 5 | 0.479 |
| 7PP2_A | 582 | 9 | 0.601 | 7VKK_A | 294 | 10 | 0.983 |
| 7T9X_A | 53 | 7 | 0.987 | 7XGB_C | 502 | 5 | 0.974 |
| 7TDV_C | 439 | 7 | 0.651 | 7XQ5_B | 67 | 10 | 0.643 |
| 7TEC_A | 82 | 7 | 0.982 | 7Y6B_A | 13 | 9 | 0.981 |
| 7F84_A | 82 | 11 | 0.967 | 7Y6C_A | 54 | 9 | 0.955 |
| 7OPK_A | 856 | 3 | 0.229 | 8CXA_A | 248 | 6 | 0.440 |
| 7PVM_A | 21 | 9 | 0.825 | 7W61_A | 242 | 7 | 0.714 |
| 7R9B_A | 221 | 8 | 0.849 | 7VQ6_A | 158 | 9 | 0.928 |
| 7RBP_A | 324 | 7 | 0.883 | 7TDR_A | 131 | 9 | 0.942 |
| 8AHP_A | 350 | 7 | 0.835 | 7SAO_A | 32 | 12 | 0.932 |
| 7U0E_H | 228 | 6 | 0.580 | 7S03_A | 107 | 9 | 0.698 |
| 7SOO_A | 84 | 12 | 0.836 | 7RMY_A | 207 | 9 | 0.990 |
| 7SNC_A | 32 | 14 | 0.981 | 7RLK_D | 116 | 10 | 0.522 |

| | | | | | | | |
|---|---|---|---|---|---|---|---|
| 7R49_B | 117 | 11 | 0.764 | 7P43_B | 678 | 6 | 0.947 |
| 7MJ3_A | 18 | 12 | 0.509 | 7P4A_A | 247 | 9 | 0.397 |
| 7LX4_A | 9 | 12 | 0.495 | 7P4N_A | 77 | 10 | 0.415 |
| 7LT7_A | 31 | 11 | 0.539 | 7P56_A | 362 | 8 | 0.140 |
| 7FH0_B | 261 | 8 | 0.965 | 7RDN_A | 204 | 10 | 0.948 |
| 7FGB_A | 387 | 8 | 0.365 | 7FAX_A | 77 | 13 | 0.964 |
| 7DZ2_C | 283 | 11 | 0.943 | 7FIA_A | 7 | 8 | 0.969 |
| 7N7G_A | 138 | 12 | 0.988 | | | | |

Table S4. Detailed information of CASP15 target
The predicted TM-score is the average of the scores of all models for the target.
The bias is obtained by averaging the absolute value of the difference between the true TM-score and the scoring for all models for this target.

| | MULTICOM_qa | | | ModFOLDdock | | |
|---|---|---|---|---|---|---|
| | Predict TM-score | Pearson | Bias | Predict TM-score | Pearson | Bias |
| H1111 | 0.478 | 0.137 | 0.659 | 0.304 | 0.239 | 0.466 |
| H1137 | 0.565 | 0.213 | 0.934 | 0.352 | 0.404 | 0.897 |
| H1114 | 0.387 | 0.012 | 0.773 | 0.374 | 0.271 | -0.097 |
| H1129 | 0.648 | 0.370 | 0.797 | 0.277 | 0.377 | 0.481 |
| H1134 | 0.844 | 0.460 | 0.982 | 0.386 | 0.585 | 0.929 |
| H1140 | 0.602 | 0.394 | 0.112 | 0.221 | 0.419 | 0.498 |
| H1141 | 0.655 | 0.416 | 0.321 | 0.239 | 0.470 | 0.258 |
| H1142 | 0.597 | 0.432 | 0.257 | 0.185 | 0.439 | 0.565 |
| H1143 | 0.799 | 0.481 | 0.959 | 0.318 | 0.553 | 0.938 |
| H1144 | 0.664 | 0.409 | 0.153 | 0.263 | 0.458 | 0.311 |
| H1151 | 0.845 | 0.430 | 0.977 | 0.414 | 0.601 | 0.950 |
| H1157 | 0.721 | 0.468 | 0.908 | 0.253 | 0.621 | 0.913 |
| H1166 | 0.699 | 0.516 | 0.856 | 0.183 | 0.531 | 0.787 |
| H1167 | 0.702 | 0.509 | 0.830 | 0.193 | 0.582 | 0.667 |
| H1168 | 0.833 | 0.577 | 0.972 | 0.257 | 0.659 | 0.918 |
| H1171 | 0.805 | 0.440 | 0.985 | 0.365 | 0.339 | -0.587 |
| H1172 | 0.783 | 0.429 | 0.990 | 0.355 | 0.343 | -0.569 |
| T1109o | 0.855 | 0.833 | 0.724 | 0.054 | 0.741 | 0.632 |
| T1110o | 0.956 | 0.850 | 0.836 | 0.108 | 0.808 | 0.849 |
| T1113o | 0.854 | 0.562 | 0.982 | 0.292 | 0.733 | 0.949 |
| T1121o | 0.564 | 0.594 | -0.242 | 0.140 | 0.655 | -0.204 |
| T1123o | 0.628 | 0.280 | 0.973 | 0.348 | 0.259 | 0.899 |
| T1124o | 0.924 | 0.694 | 0.919 | 0.230 | 0.699 | 0.893 |
| T1127o | 0.957 | 0.897 | 0.791 | 0.060 | 0.839 | 0.886 |
| T1132o | 0.893 | 0.789 | 0.930 | 0.127 | 0.648 | 0.728 |
| T1153o | 0.794 | 0.438 | 0.991 | 0.356 | 0.618 | 0.971 |
| T1161o | 0.469 | 0.740 | -0.034 | 0.282 | 0.671 | -0.330 |

| | | | | | | |
|---|---|---|---|---|---|---|
| T1170o | 0.864 | 0.502 | 0.984 | 0.363 | 0.529 | 0.558 |
| T1173o | 0.474 | 0.348 | 0.503 | 0.153 | 0.286 | 0.476 |
| T1174o | 0.629 | 0.396 | 0.852 | 0.232 | 0.392 | 0.742 |
| T1178o | 0.858 | 0.591 | 0.948 | 0.268 | 0.520 | 0.912 |
| T1179o | 0.672 | 0.357 | 0.969 | 0.314 | 0.300 | 0.899 |
| T1181o | 0.603 | 0.423 | 0.410 | 0.196 | 0.418 | 0.250 |

| | ModFOLDdockR | | | GraphGPSM | | |
|---|---|---|---|---|---|---|
| | Predict TM-score | Pearson | Bias | Predict TM-score | Pearson | Bias |
| H1111 | 0.414 | 0.447 | 0.111 | 0.341 | 0.013 | 0.185 |
| H1137 | 0.619 | 0.806 | 0.165 | 0.509 | 0.878 | 0.114 |
| H1114 | 0.461 | -0.211 | 0.285 | 0.485 | -0.602 | 0.484 |
| H1129 | 0.564 | 0.548 | 0.104 | 0.623 | 0.039 | 0.113 |
| H1134 | 0.675 | 0.851 | 0.185 | 0.836 | 0.964 | 0.040 |
| H1140 | 0.615 | 0.680 | 0.054 | 0.655 | 0.054 | 0.075 |
| H1141 | 0.625 | 0.313 | 0.056 | 0.724 | 0.283 | 0.144 |
| H1142 | 0.636 | 0.807 | 0.064 | 0.715 | 0.145 | 0.129 |
| H1143 | 0.684 | 0.673 | 0.129 | 0.817 | 0.977 | 0.024 |
| H1144 | 0.633 | 0.525 | 0.052 | 0.740 | 0.216 | 0.139 |
| H1151 | 0.686 | 0.926 | 0.168 | 0.770 | 0.953 | 0.080 |
| H1157 | 0.715 | 0.901 | 0.045 | 0.856 | 0.892 | 0.139 |
| H1166 | 0.744 | 0.767 | 0.069 | 0.751 | 0.774 | 0.066 |
| H1167 | 0.755 | 0.798 | 0.080 | 0.587 | 0.381 | 0.138 |
| H1168 | 0.774 | 0.878 | 0.078 | 0.736 | 0.799 | 0.116 |
| H1171 | 0.450 | -0.341 | 0.462 | 0.913 | 0.698 | 0.115 |
| H1172 | 0.461 | -0.339 | 0.435 | 0.876 | 0.669 | 0.104 |
| T1109o | 0.804 | 0.592 | 0.066 | 0.963 | 0.175 | 0.119 |
| T1110o | 0.828 | 0.902 | 0.130 | 0.971 | 0.654 | 0.022 |
| T1113o | 0.779 | 0.903 | 0.102 | 0.914 | 0.975 | 0.065 |
| T1121o | 0.762 | 0.034 | 0.213 | 0.406 | 0.700 | 0.192 |
| T1123o | 0.507 | 0.857 | 0.163 | 0.429 | 0.615 | 0.213 |
| T1124o | 0.805 | 0.869 | 0.121 | 0.923 | 0.967 | 0.013 |
| T1127o | 0.830 | 0.865 | 0.128 | 0.969 | 0.985 | 0.013 |
| T1132o | 0.767 | 0.782 | 0.182 | 0.831 | 0.973 | 0.068 |
| T1153o | 0.697 | 0.952 | 0.118 | 0.830 | 0.995 | 0.042 |
| T1161o | 0.767 | 0.060 | 0.303 | 0.871 | -0.411 | 0.414 |
| T1170o | 0.677 | 0.660 | 0.219 | 0.922 | 0.861 | 0.066 |
| T1173o | 0.610 | 0.399 | 0.204 | 0.545 | 0.233 | 0.190 |
| T1174o | 0.689 | 0.758 | 0.073 | 0.491 | 0.577 | 0.147 |
| T1178o | 0.700 | 0.733 | 0.180 | 0.840 | 0.973 | 0.031 |
| T1179o | 0.523 | 0.855 | 0.169 | 0.692 | 0.956 | 0.047 |
| T1181o | 0.723 | 0.377 | 0.166 | 0.573 | 0.145 | 0.187 |

| | Venclovas | | | Manifold | | |
|---|---|---|---|---|---|---|
| | Predict TM-score | Pearson | Bias | Predict TM-score | Pearson | Bias |
| H1111 | 0.436 | 0.330 | 0.225 | 0.082 | 0.206 | 0.376 |
| H1137 | 0.223 | 0.601 | 0.354 | 0.219 | 0.741 | 0.348 |
| H1114 | 0.165 | 0.593 | 0.250 | 0.195 | 0.549 | 0.220 |
| H1129 | | | | 0.517 | 0.024 | 0.147 |
| H1134 | 0.283 | 0.479 | 0.564 | 0.760 | 0.878 | 0.105 |
| H1140 | 0.062 | 0.516 | 0.554 | 0.601 | 0.048 | 0.043 |
| H1141 | 0.070 | 0.750 | 0.587 | 0.635 | 0.912 | 0.024 |
| H1142 | 0.255 | -0.102 | 0.379 | 0.658 | 0.102 | 0.098 |
| H1143 | 0.293 | 0.878 | 0.516 | 0.774 | 0.977 | 0.030 |
| H1144 | 0.149 | 0.351 | 0.530 | 0.627 | 0.188 | 0.064 |
| H1151 | 0.559 | 0.651 | 0.309 | 0.765 | 0.920 | 0.082 |
| H1157 | 0.706 | 0.828 | 0.146 | 0.727 | 0.706 | 0.105 |
| H1166 | 0.508 | 0.128 | 0.286 | 0.604 | 0.395 | 0.165 |
| H1167 | 0.467 | 0.309 | 0.277 | 0.437 | 0.159 | 0.268 |
| H1168 | 0.636 | 0.727 | 0.252 | 0.633 | 0.570 | 0.201 |
| H1171 | 0.257 | 0.086 | 0.587 | 0.610 | 0.738 | 0.201 |
| H1172 | 0.207 | 0.045 | 0.614 | 0.545 | 0.659 | 0.243 |
| T1109o | 0.772 | 0.628 | 0.105 | 0.889 | 0.668 | 0.102 |
| T1110o | 0.874 | 0.710 | 0.085 | 0.910 | 0.694 | 0.053 |
| T1113o | 0.668 | 0.693 | 0.211 | 0.841 | 0.816 | 0.080 |
| T1121o | 0.823 | -0.108 | 0.340 | 0.574 | -0.240 | 0.125 |
| T1123o | 0.168 | 0.573 | 0.464 | 0.330 | 0.842 | 0.302 |
| T1124o | 0.853 | 0.857 | 0.077 | 0.783 | 0.684 | 0.147 |
| T1127o | 0.850 | 0.746 | 0.119 | 0.928 | 0.916 | 0.036 |
| T1132o | 0.788 | 0.796 | 0.121 | 0.701 | 0.639 | 0.195 |
| T1153o | 0.631 | 0.940 | 0.198 | 0.543 | 0.811 | 0.253 |
| T1161o | 0.253 | 0.264 | 0.358 | 0.692 | 0.608 | 0.234 |
| T1170o | 0.241 | 0.259 | 0.628 | 0.274 | 0.265 | 0.592 |
| T1173o | 0.387 | 0.230 | 0.332 | 0.305 | 0.491 | 0.202 |
| T1174o | 0.513 | 0.521 | 0.307 | | | |
| T1178o | 0.570 | 0.801 | 0.296 | 0.737 | 0.930 | 0.123 |
| T1179o | 0.393 | 0.812 | 0.287 | 0.428 | 0.782 | 0.248 |
| T1181o | 0.324 | 0.075 | 0.383 | 0.325 | 0.361 | 0.290 |

| | GuijunLab-Human | | | Bhattacharya | | |
|---|---|---|---|---|---|---|
| | Predict TM-score | Pearson | Bias | Predict TM-score | Pearson | Bias |
| H1111 | | | | 0.361 | 0.209 | 0.181 |
| H1137 | 0.342 | 0.872 | 0.244 | 0.205 | 0.553 | 0.382 |
| H1114 | 0.476 | -0.590 | 0.484 | 0.340 | 0.302 | 0.202 |
| H1129 | 0.556 | 0.184 | 0.130 | 0.409 | 0.501 | 0.244 |
| H1134 | 0.713 | 0.964 | 0.134 | 0.355 | 0.668 | 0.493 |
| H1140 | 0.609 | 0.036 | 0.066 | 0.407 | 0.099 | 0.208 |
| H1141 | 0.572 | 0.146 | 0.085 | 0.433 | 0.038 | 0.223 |
| H1142 | 0.686 | 0.161 | 0.104 | 0.424 | 0.239 | 0.192 |
| H1143 | 0.610 | -0.016 | 0.216 | 0.425 | 0.404 | 0.381 |
| H1144 | 0.614 | 0.003 | 0.090 | 0.439 | 0.316 | 0.235 |
| H1151 | 0.685 | 0.952 | 0.161 | 0.327 | 0.767 | 0.520 |
| H1157 | 0.752 | 0.913 | 0.061 | 0.327 | 0.718 | 0.410 |
| H1166 | 0.673 | 0.814 | 0.066 | 0.373 | 0.661 | 0.331 |
| H1167 | 0.654 | 0.681 | 0.066 | 0.374 | 0.283 | 0.340 |
| H1168 | 0.675 | 0.881 | 0.159 | 0.403 | 0.646 | 0.443 |
| H1171 | 0.855 | 0.650 | 0.100 | 0.883 | 0.396 | 0.134 |
| H1172 | 0.834 | 0.696 | 0.101 | 0.885 | 0.453 | 0.123 |
| T1109o | 0.900 | 0.184 | 0.069 | 0.396 | 0.369 | 0.458 |
| T1110o | 0.952 | 0.644 | 0.024 | 0.399 | 0.628 | 0.556 |
| T1113o | 0.766 | 0.978 | 0.088 | 0.461 | 0.782 | 0.393 |
| T1121o | 0.609 | -0.377 | 0.179 | 0.357 | -0.152 | 0.210 |
| T1123o | 0.496 | 0.214 | 0.213 | 0.296 | 0.741 | 0.323 |
| T1124o | 0.881 | 0.948 | 0.043 | 0.327 | 0.520 | 0.602 |
| T1127o | 0.920 | 0.982 | 0.036 | 0.373 | 0.790 | 0.584 |
| T1132o | 0.800 | 0.994 | 0.093 | 0.431 | 0.593 | 0.473 |
| T1153o | 0.578 | 0.761 | 0.222 | 0.400 | 0.506 | 0.397 |
| T1161o | 0.746 | -0.293 | 0.289 | 0.290 | -0.424 | 0.183 |
| T1170o | 0.877 | 0.862 | 0.029 | 0.354 | 0.702 | 0.545 |
| T1173o | 0.524 | 0.724 | 0.113 | 0.223 | 0.381 | 0.258 |
| T1174o | 0.654 | 0.635 | 0.100 | 0.252 | 0.666 | 0.380 |
| T1178o | 0.785 | 0.994 | 0.073 | 0.362 | 0.718 | 0.504 |
| T1179o | 0.582 | 0.995 | 0.090 | 0.239 | 0.746 | 0.440 |
| T1181o | 0.576 | 0.439 | 0.161 | 0.243 | 0.242 | 0.363 |

| | VoroMQA-select-2020 | | | GuijunLab-RocketX | | |
|---|---|---|---|---|---|---|
| | Predict TM-score | Pearson | Bias | Predict TM-score | Pearson | Bias |
| H1111 | 0.498 | 0.351 | 0.246 | 0.704 | 0.082 | 0.288 |
| H1137 | 0.498 | 0.860 | 0.134 | 0.509 | 0.338 | 0.216 |
| H1114 | 0.498 | 0.545 | 0.230 | 0.617 | 0.289 | 0.383 |
| H1129 | 0.516 | 0.275 | 0.251 | 0.711 | 0.462 | 0.102 |
| H1134 | 0.498 | 0.465 | 0.371 | 0.810 | 0.639 | 0.126 |
| H1140 | 0.502 | 0.247 | 0.254 | 0.845 | 0.391 | 0.239 |
| H1141 | 0.498 | 0.192 | 0.271 | 0.881 | 0.331 | 0.232 |
| H1142 | 0.511 | 0.209 | 0.254 | 0.853 | 0.507 | 0.251 |
| H1143 | 0.502 | 0.560 | 0.342 | 0.887 | 0.517 | 0.129 |
| H1144 | 0.502 | 0.242 | 0.273 | 0.849 | 0.321 | 0.197 |
| H1151 | 0.498 | 0.496 | 0.368 | 0.745 | 0.673 | 0.156 |
| H1157 | 0.498 | 0.484 | 0.296 | | | |
| H1166 | 0.498 | 0.289 | 0.287 | 0.830 | 0.715 | 0.131 |
| H1167 | 0.498 | 0.265 | 0.294 | 0.839 | 0.732 | 0.145 |
| H1168 | 0.498 | 0.569 | 0.365 | 0.872 | 0.777 | 0.071 |
| H1171 | 0.498 | 0.518 | 0.339 | 0.883 | 0.020 | 0.148 |
| H1172 | 0.498 | 0.528 | 0.322 | 0.872 | 0.012 | 0.146 |
| T1109o | 0.498 | 0.473 | 0.367 | 0.902 | 0.354 | 0.088 |
| T1110o | 0.498 | 0.414 | 0.459 | 0.905 | 0.621 | 0.062 |
| T1113o | 0.498 | 0.581 | 0.365 | 0.872 | 0.671 | 0.064 |
| T1121o | 0.498 | -0.214 | 0.266 | 0.926 | 0.257 | 0.362 |
| T1123o | 0.498 | 0.888 | 0.155 | 0.605 | 0.744 | 0.138 |
| T1124o | 0.498 | 0.470 | 0.431 | 0.834 | 0.658 | 0.098 |
| T1127o | 0.498 | 0.440 | 0.459 | 0.897 | 0.753 | 0.064 |
| T1132o | 0.498 | 0.442 | 0.416 | 0.869 | 0.567 | 0.125 |
| T1153o | 0.498 | 0.688 | 0.313 | 0.849 | 0.320 | 0.140 |
| T1161o | 0.498 | 0.031 | 0.256 | 0.776 | 0.335 | 0.310 |
| T1170o | 0.498 | 0.530 | 0.376 | 0.882 | 0.296 | 0.074 |
| T1173o | 0.498 | 0.282 | 0.253 | 0.697 | 0.493 | 0.243 |
| T1174o | 0.498 | 0.627 | 0.233 | 0.810 | 0.766 | 0.184 |
| T1178o | 0.498 | 0.713 | 0.368 | 0.791 | 0.605 | 0.114 |
| T1179o | 0.498 | 0.635 | 0.232 | 0.636 | 0.602 | 0.143 |
| T1181o | 0.498 | 0.241 | 0.275 | 0.812 | 0.273 | 0.215 |

Table S5. Detailed information of ablation studies

| version | CASP13 | CASP14 | CAMEO |
|---|---|---|---|
| Version 1 | 0.187 | 0.191 | 0.236 |
| Version 2 | 0.238 | 0.219 | 0.275 |
| Version 3 | 0.401 | 0.382 | 0.415 |
| Version 4 | 0.458 | 0.448 | 0.518 |
| Version 5 | 0.647 | 0.69588 | 0.752 |
| Version 6 | 0.8124 | 0.7183 | 0.80949 |

Table S6. Detailed information of the 484 regular proteins

| PDB ID | TM-score of REF2015 | TM-score of REF2015+GraphGPSM |
|---|---|---|
| 1A0S_P | 0.0891 | 0.1548 |
| 1A3A_C | 0.1617 | 0.295 |
| 1A40_A | 0.12 | 0.1761 |
| 1A41_A | 0.1687 | 0.2603 |
| 1A6L_A | 0.2125 | 0.4084 |
| 1A7D_A | 0.3847 | 0.5313 |
| 1A91_A | 0.1519 | 0.2539 |
| 1ABT_A | 0.2066 | 0.3881 |
| 1ABV_A | 0.1577 | 0.2302 |
| 1AHK_A | 0.1313 | 0.2405 |
| 1AK6_A | 0.1096 | 0.2191 |
| 1AKP_A | 0.142 | 0.2243 |
| 1AOX_A | 0.1468 | 0.23 |
| 1AP7_A | 0.167 | 0.3786 |
| 1AUU_A | 0.1827 | 0.3046 |
| 1AX8_A | 0.141 | 0.2131 |
| 1AZ0_A | 0.2112 | 0.417 |
| 1B4B_A | 0.1664 | 0.2766 |
| 1B4R_A | 0.183 | 0.3907 |
| 1B4U_A | 0.1597 | 0.2328 |
| 1B8Q_A | 0.4029 | 0.526 |
| 1BE3_J | 0.2357 | 0.3988 |
| 1BG8_A | 0.2003 | 0.3997 |
| 1BGF_A | 0.3642 | 0.537 |
| 1BGY_J | 0.1525 | 0.2839 |
| 1BJX_A | 0.1625 | 0.2682 |
| 1BUO_A | 0.1817 | 0.3013 |
| 1C03_A | 0.1439 | 0.2349 |
| 1C1K_A | 0.2214 | 0.2628 |
| 1C41_A | 0.1802 | 0.2398 |
| 1C5E_A | 0.248 | 0.2823 |
| 1C9F_A | 0.1797 | 0.2465 |

| | | |
|---|---|---|
| 1CDB_A | 0.2147 | 0.4316 |
| 1CF7_B | 0.1931 | 0.2592 |
| 1CHC_A | 0.208 | 0.2853 |
| 1CQA_A | 0.1332 | 0.2008 |
| 1CTO_A | 0.3187 | 0.5028 |
| 1CXZ_B | 0.2039 | 0.3093 |
| 1D6T_A | 0.2592 | 0.356 |
| 1D8B_A | 0.1698 | 0.2858 |
| 1DBF_A | 0.1983 | 0.3825 |
| 1DCF_A | 0.1837 | 0.3497 |
| 1DJ7_A | 0.1817 | 0.3421 |
| 1DL6_A | 0.1434 | 0.2556 |
| 1DOI_A | 0.2843 | 0.4465 |
| 1DP7_A | 0.2014 | 0.5071 |
| 1DP7_P | 0.1449 | 0.2012 |
| 1DTP_A | 0.1522 | 0.2491 |
| 1DTV_A | 0.1064 | 0.2443 |
| 1DUN_A | 0.2283 | 0.3798 |
| 1DWM_A | 0.3353 | 0.6241 |
| 1E2A_A | 0.195 | 0.3458 |
| 1E3Y_A | 0.2107 | 0.3118 |
| 1E53_A | 0.1578 | 0.2405 |
| 1EGG_B | 0.2204 | 0.3403 |
| 1EKZ_A | 0.1709 | 0.491 |
| 1ELW_A | 0.1853 | 0.2961 |
| 1EM8_D | 0.1751 | 0.4351 |
| 1EZV_G | 0.1156 | 0.1972 |
| 1F15_C | 0.2031 | 0.329 |
| 1F1E_A | 0.1695 | 0.3279 |
| 1F2R_I | 0.2038 | 0.2871 |
| 1F37_B | 0.2001 | 0.3221 |
| 1F3U_A | 0.1367 | 0.1959 |
| 1F3Y_A | 0.2433 | 0.6009 |
| 1F43_A | 0.1852 | 0.2673 |
| 1F5X_A | 0.2056 | 0.367 |
| 1F93_A | 0.2079 | 0.3762 |
| 1F98_A | 0.1889 | 0.3892 |
| 1F9P_A | 0.1355 | 0.2863 |
| 1FAQ_A | 0.1776 | 0.352 |
| 1FC3_A | 0.1529 | 0.395 |
| 1FCA_A | 0.2272 | 0.487 |
| 1FEX_A | 0.1571 | 0.2975 |
| 1FHT_A | 0.3414 | 0.3657 |
| 1FJG_F | 0.1297 | 0.2112 |

| | | |
|---|---|---|
| 1FJG_H | 0.1288 | 0.3052 |
| 1FMB_A | 0.1984 | 0.3532 |
| 1FR0_A | 0.151 | 0.3268 |
| 1FRD_A | 0.3775 | 0.3649 |
| 1FSP_A | 0.1063 | 0.1478 |
| 1FSU_A | 0.1382 | 0.1918 |
| 1FW9_A | 0.1071 | 0.2108 |
| 1FY2_A | 0.2455 | 0.3589 |
| 1G2R_A | 0.2574 | 0.3742 |
| 1G8Q_A | 0.2094 | 0.4638 |
| 1GGS_A | 0.1384 | 0.2267 |
| 1GME_A | 0.1722 | 0.2485 |
| 1GPQ_B | 0.2029 | 0.3724 |
| 1GQA_A | 0.2854 | 0.4477 |
| 1GVN_A | 0.1715 | 0.3608 |
| 1GVP_A | 0.1243 | 0.2186 |
| 1GXD_C | 0.1808 | 0.2847 |
| 1GXL_A | 0.1254 | 0.1636 |
| 1GYH_A | 0.1777 | 0.2951 |
| 1H4L_D | 0.2142 | 0.2863 |
| 1H8E_H | 0.1965 | 0.3659 |
| 1H9E_A | 0.2105 | 0.3823 |
| 1H9F_A | 0.1747 | 0.36 |
| 1HBG_A | 0.2524 | 0.4227 |
| 1HBX_E | 0.2264 | 0.2545 |
| 1HCD_A | 0.1654 | 0.2227 |
| 1HH8_A | 0.1539 | 0.2901 |
| 1HHV_A | 0.1813 | 0.3593 |
| 1HKQ_A | 0.1782 | 0.3293 |
| 1HKX_E | 0.1521 | 0.2616 |
| 1HL6_D | 0.2236 | 0.436 |
| 1I1N_A | 0.1561 | 0.2815 |
| 1I27_A | 0.2283 | 0.2663 |
| 1I35_A | 0.163 | 0.231 |
| 1I85_A | 0.1239 | 0.216 |
| 1ID2_A | 0.1403 | 0.3186 |
| 1IEZ_A | 0.155 | 0.2653 |
| 1IJY_A | 0.1501 | 0.233 |
| 1IM3_D | 0.182 | 0.3185 |
| 1IOO_A | 0.2026 | 0.3628 |
| 1IPI_A | 0.1869 | 0.3174 |
| 1IQV_A | 0.1463 | 0.3261 |
| 1IRS_A | 0.1992 | 0.3705 |
| 1IS7_K | 0.2154 | 0.3216 |

| | | |
|---|---|---|
| 1IUJ_B | 0.3238 | 0.4044 |
| 1IUY_A | 0.2029 | 0.3239 |
| 1J1V_A | 0.1743 | 0.3222 |
| 1J3W_A | 0.2293 | 0.4272 |
| 1J8I_A | 0.1494 | 0.2074 |
| 1J9I_A | 0.3093 | 0.4142 |
| 1JC7_A | 0.1671 | 0.2628 |
| 1JEI_A | 0.1388 | 0.2378 |
| 1JIW_I | 0.1543 | 0.2864 |
| 1JJ2_S | 0.2262 | 0.3934 |
| 1JLI_A | 0.1924 | 0.3082 |
| 1JMT_A | 0.1221 | 0.1875 |
| 1JO0_A | 0.164 | 0.2622 |
| 1JOF_A | 0.1521 | 0.2381 |
| 1JOP_A | 0.241 | 0.3527 |
| 1JPY_Y | 0.2118 | 0.4178 |
| 1JR5_A | 0.139 | 0.2175 |
| 1JR8_A | 0.135 | 0.3086 |
| 1JSG_A | 0.1378 | 0.2592 |
| 1K1Z_A | 0.188 | 0.3502 |
| 1K3B_A | 0.1591 | 0.2463 |
| 1K3S_A | 0.2436 | 0.3138 |
| 1K5D_B | 0.2131 | 0.3577 |
| 1K73_1 | 0.1777 | 0.3685 |
| 1KA8_A | 0.1334 | 0.248 |
| 1KN6_A | 0.1603 | 0.2574 |
| 1KOH_D | 0.1587 | 0.2707 |
| 1KP6_A | 0.1965 | 0.3581 |
| 1KPT_A | 0.1735 | 0.2728 |
| 1KQ6_A | 0.2023 | 0.3557 |
| 1KSX_A | 0.2708 | 0.4576 |
| 1KVD_B | 0.1236 | 0.2084 |
| 1KX5_D | 0.3993 | 0.9327 |
| 1L1D_B | 0.1602 | 0.3002 |
| 1L2P_A | 0.1862 | 0.3634 |
| 1L3G_A | 0.2526 | 0.4109 |
| 1L6H_A | 0.1784 | 0.3279 |
| 1LDD_A | 0.2411 | 0.4539 |
| 1LE2_A | 0.1343 | 0.2057 |
| 1LFU_P | 0.2909 | 0.3572 |
| 1LMI_A | 0.1756 | 0.2567 |
| 1LNW_C | 0.3439 | 0.5569 |
| 1LQM_H | 0.166 | 0.3301 |
| 1LR1_B | 0.1258 | 0.2218 |

| | | |
|---|---|---|
| 1LWB_A | 0.1997 | 0.3354 |
| 1LYV_A | 0.1079 | 0.1641 |
| 1LZW_B | 0.1657 | 0.2536 |
| 1M7Y_A | 0.1379 | 0.3446 |
| 1MAI_A | 0.2547 | 0.43 |
| 1MC2_A | 0.1506 | 0.2863 |
| 1MFQ_C | 0.0828 | 0.156 |
| 1MHD_A | 0.1729 | 0.2843 |
| 1MKF_A | 0.0894 | 0.2219 |
| 1MN8_A | 0.2044 | 0.3141 |
| 1MT3_A | 0.219 | 0.3068 |
| 1MWP_A | 0.1361 | 0.1956 |
| 1MWQ_A | 0.2151 | 0.293 |
| 1N12_A | 0.2292 | 0.4604 |
| 1N13_B | 0.2398 | 0.337 |
| 1N3G_A | 0.1895 | 0.3543 |
| 1N8V_A | 0.1933 | 0.258 |
| 1NF6_F | 0.3763 | 0.6281 |
| 1NGL_A | 0.1347 | 0.226 |
| 1NKZ_A | 0.1555 | 0.2541 |
| 1NLQ_A | 0.1737 | 0.3109 |
| 1NOE_A | 0.2313 | 0.3212 |
| 1NPB_A | 0.1282 | 0.1848 |
| 1NQJ_B | 0.1897 | 0.2553 |
| 1NQZ_A | 0.1866 | 0.214 |
| 1NR3_A | 0.196 | 0.2761 |
| 1NRJ_B | 0.2628 | 0.4553 |
| 1NTV_A | 0.1632 | 0.2474 |
| 1NZE_A | 0.1617 | 0.2193 |
| 1O0G_A | 0.2112 | 0.2687 |
| 1O9G_A | 0.1616 | 0.2725 |
| 1OA8_D | 0.19 | 0.2602 |
| 1OFT_A | 0.138 | 0.2922 |
| 1OJG_A | 0.1567 | 0.3138 |
| 1OK0_A | 0.1057 | 0.176 |
| 1OOF_A | 0.1967 | 0.2733 |
| 1OPO_C | 0.2245 | 0.3959 |
| 1ORQ_C | 0.1729 | 0.2693 |
| 1ORY_A | 0.1985 | 0.3426 |
| 1OX7_A | 0.1318 | 0.2055 |
| 1OZ9_A | 0.1895 | 0.2416 |
| 1P9O_B | 0.1272 | 0.2472 |
| 1PD6_A | 0.1347 | 0.2357 |
| 1PFS_A | 0.1672 | 0.2672 |

| | | |
|---|---|---|
| 1PGV_A | 0.1586 | 0.2817 |
| 1PIH_A | 0.233 | 0.4417 |
| 1PMS_A | 0.1678 | 0.2741 |
| 1PSR_A | 0.3378 | 0.415 |
| 1PXW_A | 0.1659 | 0.2371 |
| 1PZW_A | 0.2028 | 0.2286 |
| 1QFT_A | 0.1709 | 0.2587 |
| 1QFW_B | 0.1229 | 0.2006 |
| 1QMA_A | 0.1162 | 0.2102 |
| 1QWT_A | 0.1671 | 0.2791 |
| 1QZG_A | 0.1184 | 0.2028 |
| 1R5T_A | 0.3356 | 0.4722 |
| 1R5Z_A | 0.1547 | 0.2591 |
| 1R6R_A | 0.1213 | 0.1653 |
| 1RHX_A | 0.1764 | 0.2274 |
| 1RKX_A | 0.2352 | 0.2947 |
| 1ROW_A | 0.1259 | 0.2044 |
| 1RTU_A | 0.1373 | 0.2739 |
| 1RZ2_A | 0.1591 | 0.2627 |
| 1RZ3_A | 0.2258 | 0.3961 |
| 1S2D_A | 0.1964 | 0.3276 |
| 1S3J_A | 0.2351 | 0.4784 |
| 1S56_B | 0.2264 | 0.3472 |
| 1S7O_C | 0.2072 | 0.3406 |
| 1S7Z_A | 0.1347 | 0.2417 |
| 1SAU_A | 0.1752 | 0.2843 |
| 1SG4_B | 0.129 | 0.2086 |
| 1SMP_I | 0.1359 | 0.1722 |
| 1SPP_B | 0.1567 | 0.1867 |
| 1SR8_A | 0.1473 | 0.2623 |
| 1STM_A | 0.4141 | 0.6599 |
| 1SVJ_A | 0.1732 | 0.2313 |
| 1TAF_A | 0.1615 | 0.3056 |
| 1TEO_A | 0.2012 | 0.2476 |
| 1TJF_B | 0.1548 | 0.2671 |
| 1TLJ_A | 0.1346 | 0.2709 |
| 1TUL_A | 0.1888 | 0.3301 |
| 1TWU_A | 0.2407 | 0.2833 |

| | | |
|---|---|---|
| 1TYG_B | 0.2328 | 0.4875 |
| 1TZ0_A | 0.1818 | 0.2937 |
| 1U84_A | 0.2377 | 0.4273 |
| 1UDD_A | 0.179 | 0.258 |
| 1UFB_A | 0.1619 | 0.257 |
| 1UG4_A | 0.1544 | 0.2736 |
| 1UGI_A | 0.1284 | 0.2438 |
| 1UNG_D | 0.2172 | 0.4044 |
| 1USL_C | 0.1772 | 0.3115 |
| 1V74_A | 0.1414 | 0.2261 |
| 1VCC_A | 0.115 | 0.2118 |
| 1VCY_A | 0.0973 | 0.1828 |
| 1VD0_A | 0.1207 | 0.2232 |
| 1VDW_A | 0.194 | 0.3574 |
| 1VHG_A | 0.1316 | 0.2026 |
| 1VKE_E | 0.1201 | 0.1903 |
| 1VLL_A | 0.1898 | 0.3271 |
| 1VTK_A | 0.1894 | 0.2924 |
| 1VYI_A | 0.2652 | 0.3969 |
| 1VYX_A | 0.2649 | 0.5271 |
| 1W1W_E | 0.1232 | 0.2209 |
| 1W53_A | 0.2791 | 0.4079 |
| 1WER_A | 0.0822 | 0.1551 |
| 1WJ8_A | 0.1738 | 0.2953 |
| 1WKR_A | 0.1825 | 0.3393 |
| 1WLQ_C | 0.1581 | 0.2492 |
| 1WMH_B | 0.1734 | 0.312 |
| 1XJA_C | 0.1525 | 0.313 |
| 1Y14_A | 0.2202 | 0.3319 |
| 1Y1X_A | 0.1278 | 0.2034 |
| 1YG2_A | 0.1077 | 0.2005 |
| 1YZH_B | 0.1091 | 0.1688 |
| 1Z8R_A | 0.1569 | 0.2846 |
| 1ZC9_A | 0.2908 | 0.3999 |
| 2A5Y_A | 0.1346 | 0.2047 |
| 2A9U_B | 0.1924 | 0.4507 |
| 2AAM_B | 0.1606 | 0.1931 |
| 2ACY_A | 0.1523 | 0.2663 |
| 2AEN_A | 0.2323 | 0.4367 |
| 2APN_A | 0.1263 | 0.1976 |
| 2AQ0_A | 0.1362 | 0.2025 |
| 2AQS_A | 0.2464 | 0.4226 |
| 2AQX_A | 0.1311 | 0.2501 |

| | | |
|---|---|---|
| 2BL7_A | 0.2466 | 0.315 |
| 2BS2_C | 0.1469 | 0.2824 |
| 2BSE_A | 0.1519 | 0.2709 |
| 2BT9_A | 0.1571 | 0.2536 |
| 2BTD_A | 0.3289 | 0.5518 |
| 2BWJ_A | 0.201 | 0.35 |
| 2BYK_D | 0.13 | 0.2445 |
| 2C2F_A | 0.1753 | 0.2446 |
| 2C4W_A | 0.1578 | 0.232 |
| 2CDP_A | 0.287 | 0.4016 |
| 2CEY_A | 0.1616 | 0.2369 |
| 2CMX_A | 0.1361 | 0.2578 |
| 2CO3_B | 0.1671 | 0.3106 |
| 2CWP_A | 0.1616 | 0.3141 |
| 2CZV_D | 0.2148 | 0.3198 |
| 2D0P_B | 0.0974 | 0.1926 |
| 2D5R_B | 0.1954 | 0.3256 |
| 2DDS_A | 0.2097 | 0.3076 |
| 2EWC_B | 0.2525 | 0.3943 |
| 2F22_B | 0.1174 | 0.246 |
| 2FA5_B | 0.1725 | 0.3495 |
| 2FKB_C | 0.1782 | 0.3058 |
| 2GBJ_B | 0.2006 | 0.2673 |
| 2GJ3_A | 0.1455 | 0.2529 |
| 2GKC_A | 0.1777 | 0.2773 |
| 2H30_A | 0.2511 | 0.5051 |
| 2H8E_A | 0.1441 | 0.2879 |
| 2HI3_A | 0.1622 | 0.2767 |
| 2HQ7_B | 0.23 | 0.4799 |
| 2HYB_A | 0.1392 | 0.1973 |
| 2ICT_A | 0.1296 | 0.2308 |
| 2IEE_A | 0.1899 | 0.3085 |
| 2IGS_B | 0.1342 | 0.204 |
| 2J4B_B | 0.1254 | 0.2591 |
| 2J4H_B | 0.309 | 0.4926 |
| 2J5S_A | 0.1244 | 0.2047 |
| 2J6Z_A | 0.1158 | 0.1932 |
| 2JC5_A | 0.3779 | 0.4516 |
| 2JLP_D | 0.1538 | 0.2724 |
| 2JP3_A | 0.1373 | 0.3271 |
| 2K9X_A | 0.1855 | 0.3342 |
| 2KBW_A | 0.157 | 0.2998 |
| 2L5P_A | 0.2298 | 0.3646 |

| | | |
|---|---|---|
| 2L74_A | 0.1575 | 0.2486 |
| 2LKP_A | 0.1569 | 0.2895 |
| 2LRB_A | 0.1533 | 0.3021 |
| 2LWP_A | 0.2172 | 0.406 |
| 2NAZ_A | 0.2309 | 0.4365 |
| 2NCM_A | 0.1874 | 0.3051 |
| 2NDP_A | 0.1016 | 0.1834 |
| 2NS9_B | 0.1373 | 0.1944 |
| 2NZY_A | 0.1898 | 0.3035 |
| 2O2Y_C | 0.3307 | 0.6384 |
| 2O70_F | 0.1625 | 0.2736 |
| 2ODM_B | 0.1827 | 0.2899 |
| 2P7L_A | 0.1293 | 0.1862 |
| 2PI2_F | 0.1146 | 0.1973 |
| 2PK3_A | 0.2405 | 0.3753 |
| 2PPV_A | 0.1288 | 0.2133 |
| 2PYB_A | 0.1431 | 0.229 |
| 2Q2H_A | 0.1513 | 0.3638 |
| 2QTT_A | 0.1874 | 0.4775 |
| 2QVG_A | 0.1312 | 0.186 |
| 2QZJ_A | 0.1321 | 0.2531 |
| 2R8W_B | 0.186 | 0.2685 |
| 2RCC_B | 0.2867 | 0.4429 |
| 2RD5_D | 0.1588 | 0.2538 |
| 2RLD_C | 0.1556 | 0.3152 |
| 2UUX_A | 0.1117 | 0.1655 |
| 2V85_A | 0.1399 | 0.2241 |
| 2VDX_A | 0.1637 | 0.2894 |
| 2VUL_A | 0.1444 | 0.2584 |
| 2WCW_B | 0.1002 | 0.1664 |
| 2WGP_A | 0.1826 | 0.3303 |
| 2WZR_1 | 0.112 | 0.1737 |
| 2XGY_A | 0.1845 | 0.2669 |
| 2XW2_A | 0.186 | 0.2757 |
| 2Z3B_A | 0.108 | 0.1759 |
| 2ZMZ_B | 0.0947 | 0.152 |
| 3A04_A | 0.1197 | 0.2263 |
| 3A3K_B | 0.1559 | 0.2388 |
| 3ALU_A | 0.1751 | 0.2741 |
| 3BDB_A | 0.2462 | 0.4334 |
| 3CAE_A | 0.2014 | 0.3015 |
| 3CG4_A | 0.2971 | 0.5372 |
| 3CHB_D | 0.2316 | 0.3555 |

| | | |
|---|---|---|
| 3CX5_F | 0.2511 | 0.3467 |
| 3CX5_G | 0.2617 | 0.2001 |
| 3E6M_E | 0.2841 | 0.3654 |
| 3E9T_D | 0.1738 | 0.2496 |
| 3EOD_A | 0.2059 | 0.2634 |
| 3F4W_A | 0.2349 | 0.3512 |
| 3F8L_A | 0.1107 | 0.2114 |
| 3G20_B | 0.1514 | 0.261 |
| 3GMX_A | 0.1502 | 0.2173 |
| 3H05_B | 0.1989 | 0.2704 |
| 3HGI_A | 0.1528 | 0.2386 |
| 3I9V_7 | 0.1378 | 0.1864 |
| 3IAM_2 | 0.1092 | 0.2031 |
| 3IAM_4 | 0.2106 | 0.3324 |
| 3JB9_L | 0.1303 | 0.1988 |
| 3JBR_E | 0.1114 | 0.1632 |
| 3JU7_A | 0.2143 | 0.2855 |
| 3JZ4_A | 0.1358 | 0.2376 |
| 3LQV_B | 0.1403 | 0.2365 |
| 3M1N_B | 0.1262 | 0.2897 |
| 3MAT_A | 0.1022 | 0.1651 |
| 3MQK_C | 0.0995 | 0.1939 |
| 3MX3_A | 0.2159 | 0.2862 |
| 3N0X_A | 0.2197 | 0.2993 |
| 3N1G_C | 0.122 | 0.2057 |
| 3N9U_C | 0.1767 | 0.2112 |
| 3NGF_A | 0.1151 | 0.1934 |
| 3O61_A | 0.173 | 0.2825 |
| 3OUJ_A | 0.1728 | 0.2647 |
| 3P8B_A | 0.1038 | 0.2052 |
| 3PD2_A | 0.0931 | 0.1748 |
| 3QFM_A | 0.1742 | 0.2727 |
| 3QNS_A | 0.1249 | 0.2115 |
| 3QU3_A | 0.1298 | 0.2098 |
| 3R9T_A | 0.2047 | 0.3422 |
| 3ROT_A | 0.1456 | 0.2439 |
| 3SDL_B | 0.2182 | 0.3852 |
| 3TEO_A | 0.1378 | 0.2544 |
| 3UE6_E | 0.1557 | 0.2678 |
| 3V1O_A | 0.1924 | 0.3855 |
| 3W1Z_D | 0.2043 | 0.3627 |
| 3X0G_A | 0.129 | 0.2019 |
| 3X15_A | 0.153 | 0.2264 |

| | | |
|---|---|---|
| 3ZGF_A | 0.2533 | 0.4104 |
| 3ZO5_A | 0.1804 | 0.326 |
| 4AIH_A | 0.1063 | 0.1837 |
| 4ASW_C | 0.1158 | 0.2703 |
| 4AUB_C | 0.1048 | 0.1751 |
| 4B0M_A | 0.1979 | 0.2695 |
| 4CSD_A | 0.094 | 0.1935 |
| 4CXT_A | 0.1469 | 0.2421 |
| 4DF3_A | 0.2575 | 0.4032 |
| 4DYW_A | 0.1297 | 0.201 |
| 4ESB_A | 0.1718 | 0.3487 |
| 4F8X_A | 0.1568 | 0.299 |
| 4GDK_A | 0.1431 | 0.2568 |
| 4GF3_A | 0.1504 | 0.3649 |
| 4GQY_A | 0.0871 | 0.1725 |
| 4I60_A | 0.1258 | 0.2485 |
| 4IMH_A | 0.186 | 0.3201 |
| 4IOS_A | 0.2094 | 0.3215 |
| 4J20_A | 0.101 | 0.2175 |
| 4JGX_B | 0.1622 | 0.2565 |
| 4K1F_A | 0.0996 | 0.2047 |
| 4KA0_A | 0.1184 | 0.1822 |
| 4KCD_B | 0.1518 | 0.4121 |
| 4KKY_X | 0.2421 | 0.3491 |
| 4LE0_B | 0.0965 | 0.2007 |
| 4LMS_A | 0.156 | 0.3334 |
| 4LZ6_A | 0.1378 | 0.2365 |
| 4M75_F | 0.1929 | 0.2666 |
| 4M7O_A | 0.1893 | 0.3368 |
| 4MLF_D | 0.1159 | 0.1914 |
| 4MMG_A | 0.1699 | 0.2438 |
| 4MOU_A | 0.2587 | 0.3676 |
| 4NBI_A | 0.1653 | 0.2825 |
| 4OW1_A | 0.1882 | 0.2873 |
| 4Q2O_A | 0.1129 | 0.1752 |
| 4Q2Q_A | 0.1541 | 0.2455 |
| 4Q75_A | 0.1172 | 0.1753 |
| 4R67_0 | 0.2408 | 0.3596 |
| 4R8D_A | 0.1866 | 0.3005 |
| 4RUV_A | 0.2388 | 0.5302 |
| 4UIJ_A | 0.1447 | 0.221 |
| 4V2O_A | 0.1697 | 0.2461 |
| 4XEQ_B | 0.1326 | 0.22 |

| | | |
|---|---|---|
| 4Z6J_A | 0.1243 | 0.1759 |
| 4ZBY_A | 0.1053 | 0.1837 |
| 4ZDO_A | 0.1498 | 0.2946 |
| 4ZF6_A | 0.1275 | 0.2427 |
| 5CJ3_B | 0.176 | 0.332 |
| 5CZ8_Y | 0.1252 | 0.2242 |
| 5E4E_A | 0.1378 | 0.211 |
| 5EKT_A | 0.1681 | 0.3259 |
| 5IAO_A | 0.2832 | 0.262 |
| 5IZB_A | 0.2221 | 0.3582 |
| 5JTM_A | 0.1199 | 0.2422 |
| 5L38_A | 0.1194 | 0.2427 |
| 5L8R_D | 0.107 | 0.1832 |
| 5LAI_L | 0.1961 | 0.4302 |
| 5LXE_A | 0.1952 | 0.2819 |
| 5O2V_A | 0.2141 | 0.3492 |
| 5O8G_A | 0.1766 | 0.3288 |
| 5T17_A | 0.2727 | 0.4799 |
| 5TMF_E | 0.134 | 0.2039 |
| 5TUV_B | 0.1164 | 0.1757 |
| 5VAZ_B | 0.1543 | 0.2426 |
| 5W71_A | 0.1625 | 0.2998 |
| 5WSE_A | 0.1582 | 0.2135 |

Table S7. Detailed information of the 35 orphan proteins

| PDB ID | TM-score of AlphaFold2 | TM-score of GraphGPSM |
|---|---|---|
| 7A8S_1 | 0.6086 | 0.6293 |
| 7AD5_1 | 0.181 | 0.2638 |
| 7AH0_1 | 0.4355 | 0.4675 |
| 7BNY_1 | 0.6417 | 0.6527 |
| 7CG5_1 | 0.1088 | 0.2146 |
| 7DGW_1 | 0.9018 | 0.9039 |
| 7DNS_1 | 0.3843 | 0.404 |
| 7K3H_1 | 0.2182 | 0.3097 |
| 7KUW_1 | 0.6569 | 0.704 |
| 7KWW_1 | 0.3939 | 0.3939 |
| 7MQQ_1 | 0.154 | 0.1743 |
| 7S5L_1 | 0.9802 | 0.9802 |
| 7TJL_1 | 0.4916 | 0.531 |
| 7WRK_1 | 0.6977 | 0.1806 |
| 1Q2I_A | 0.34676 | 0.45953 |
| 1UVQ_C | 0.26268 | 0.33947 |

| | | |
|---|---|---|
| 2DCO_A | 0.32944 | 0.4245 |
| 2KWZ_A | 0.48608 | 0.61722 |
| 3C0T_B | 0.88776 | 0.88776 |
| 3UL1_A | 0.14873 | 0.28134 |
| 5UOI_A | 0.81632 | 0.81632 |
| 5UP5_A | 0.70319 | 0.70319 |
| 6E5N_A | 0.56594 | 0.64704 |
| 7AL0_A | 0.78159 | 0.79026 |
| 1C94_A | 0.84476 | 0.84476 |
| 2Z3F_I | 0.33226 | 0.44485 |
| 3NK3_C | 0.24917 | 0.33309 |
| 3SRJ_C | 0.19505 | 0.19607 |
| 3TWE_A | 0.81767 | 0.82388 |
| 3UKW_C | 0.23692 | 0.36946 |
| 4H25_C | 0.23166 | 0.30329 |
| 4J4A_A | 0.87668 | 0.87668 |
| 4JWE_C | 0.33721 | 0.613 |
| 4TZS_C | 0.22528 | 0.26398 |
| 4WFD_C | 0.3086 | 0.566 |
| 5JGE_C | 0.59746 | 0.69565 |

Table S8. Detailed information of the 57 multi-domain proteins

| PDB ID | TM-score of AlphaFold2 | TM-score of GraphGPSM |
|---|---|---|
| 1B3U_1 | 0.72 | 0.77 |
| 1GGZ_1 | 0.59 | 0.63 |
| 1GRI_1 | 0.46 | 0.46 |
| 1J1J_1 | 0.97 | 0.97 |
| 1Q8K_1 | 0.6 | 0.55 |
| 1QBK_1 | 0.79 | 0.83 |
| 1S8O_1 | 0.72 | 0.72 |
| 1ST0_1 | 0.78 | 0.78 |
| 1XA6_1 | 0.54 | 0.56 |
| 1YTQ_1 | 0.49 | 0.27 |
| 1ZZA_1 | 0.34 | 0.52 |
| 216L_1 | 0.38 | 0.37 |
| 2AR7_1 | 0.75 | 0.74 |
| 2DYB_1 | 0.71 | 0.74 |
| 2EYZ_1 | 0.32 | 0.24 |
| 2F83_1 | 0.69 | 0.71 |
| 2GF5_1 | 0.48 | 0.49 |
| 2KDO_1 | 0.43 | 0.34 |
| 2KN6_1 | 0.45 | 0.47 |
| 2KR0_1 | 0.31 | 0.34 |

| | | |
|---|---|---|
| 2KTY_1 | 0.8 | 0.78 |
| 2L4H_1 | 0.75 | 0.73 |
| 2LOO_1 | 0.36 | 0.44 |
| 2MPH_1 | 0.5 | 0.27 |
| 2MZH_1 | 0.74 | 0.75 |
| 2OK5_1 | 0.59 | 0.6 |
| 2P01_1 | 0.28 | 0.43 |
| 2WZB_1 | 0.77 | 0.78 |
| 2YMB_1 | 0.67 | 0.74 |
| 3AJM_1 | 0.69 | 0.71 |
| 3AY5_1 | 0.98 | 0.98 |
| 3J0A_1 | 0.62 | 0.67 |
| 4A5T_1 | 0.72 | 0.62 |
| 4BIT_1 | 0.53 | 0.62 |
| 4COO_1 | 0.75 | 0.75 |
| 4D1E_1 | 0.42 | 0.48 |
| 4IGG_1 | 0.67 | 0.67 |
| 4KFZ_1 | 0.5 | 0.49 |
| 4MSP_1 | 0.67 | 0.72 |
| 5C19_1 | 0.78 | 0.64 |
| 5EDM_1 | 0.73 | 0.69 |
| 5IVW_1 | 0.4 | 0.36 |
| 6D6Q_1 | 0.72 | 0.72 |
| 6FON_1 | 0.62 | 0.68 |
| 6HHJ_1 | 0.67 | 0.66 |
| 6I7S_1 | 0.61 | 0.6 |
| 6JT0_2 | 0.45 | 0.52 |
| 6KVG_1 | 0.51 | 0.51 |
| 6NR8_1 | 0.7 | 0.51 |
| 6R6H_2 | 0.73 | 0.75 |
| 6SN1_2 | 0.73 | 0.79 |
| 6WM2_8 | 0.7 | 0.33 |
| 6Y4L_1 | 0.75 | 0.78 |
| 7A09_10 | 0.77 | 0.43 |
| 7CCC_2 | 0.6 | 0.6 |
| 7DME_1 | 0.43 | 0.27 |

Table S9. Comparison of the model selected by GraphGPSM and the first model of AlphaFold2 on the 66 targets of CASP14. The table lists the TM-scores of the first model of AlphaFold2, the model chosen by GraphGPSM, and the best model among the top-5 models, respectively. The numbers in parentheses indicate the index of the selected model.

| Target | First model | GraphGPSM selected model | Best model | Target | First model | GraphGPSM selected model | Best model |
|---|---|---|---|---|---|---|---|
| T1024 | 0.671 (1) | 0.930 (3) | 0.930 (3) | T1060s3 | 0.957 (1) | 0.952 (3) | 0.957 (1) |
| T1026 | 0.957 (1) | 0.957 (1) | 0.957 (1) | T1061 | 0.768 (1) | 0.778 (4) | 0.792 (3) |
| T1027 | 0.485 (1) | 0.483 (2) | 0.488 (4) | T1062 | 0.571 (1) | 0.574 (4) | 0.574 (4) |
| T1029 | 0.469 (1) | 0.471 (4) | 0.471 (4) | T1064 | 0.812 (1) | 0.812 (1) | 0.812 (1) |
| T1030 | 0.730 (1) | 0.740 (2) | 0.740 (2) | T1065s1 | 0.958 (1) | 0.957 (4) | 0.962 (2) |
| T1031 | 0.855 (1) | 0.854 (2) | 0.870 (4) | T1065s2 | 0.976 (1) | 0.976 (5) | 0.976 (1) |
| T1032 | 0.707 (1) | 0.702 (4) | 0.708 (2) | T1067 | 0.925 (1) | 0.930 (5) | 0.930 (3) |
| T1033 | 0.889 (1) | 0.878 (5) | 0.889 (1) | T1068 | 0.971 (1) | 0.973 (4) | 0.973 (2) |
| T1034 | 0.959 (1) | 0.959 (1) | 0.959 (1) | T1070 | 0.488 (1) | 0.529 (5) | 0.488 (1) |
| T1035 | 0.946 (1) | 0.957 (5) | 0.957 (5) | T1072s1 | 0.749 (1) | 0.744 (3) | 0.749 (1) |
| T1037 | 0.959 (1) | 0.975 (2) | 0.975 (4) | T1073 | 0.768 (1) | 0.769 (2) | 0.769 (3) |
| T1038 | 0.917 (1) | 0.919 (5) | 0.919 (2) | T1074 | 0.916 (1) | 0.916 (3) | 0.925 (3) |
| T1039 | 0.861 (1) | 0.861 (1) | 0.861 (1) | T1076 | 0.995 (1) | 0.995 (2) | 0.995 (3) |
| T1040 | 0.784 (1) | 0.784 (1) | 0.784 (1) | T1078 | 0.951 (1) | 0.956 (1) | 0.961 (3) |
| T1041 | 0.953 (1) | 0.951 (2) | 0.953 (1) | T1079 | 0.983 (1) | 0.983 (4) | 0.983 (2) |
| T1042 | 0.936 (1) | 0.949 (3) | 0.949 (3) | T1080 | 0.876 (1) | 0.887 (3) | 0.895 (4) |
| T1043 | 0.879 (1) | 0.884 (3) | 0.884 (3) | T1082 | 0.934 (1) | 0.935 (4) | 0.935 (4) |
| T1045s2 | 0.941 (1) | 0.949 (3) | 0.941 (1) | T1083 | 0.833 (1) | 0.837 (2) | 0.877 (4) |
| T1046s1 | 0.955 (1) | 0.955 (3) | 0.956 (4) | T1084 | 0.885 (1) | 0.902 (5) | 0.909 (2) |
| T1046s2 | 0.978 (1) | 0.978 (1) | 0.978 (1) | T1087 | 0.952 (1) | 0.946 (5) | 0.952 (1) |
| T1047s1 | 0.559 (1) | 0.559 (1) | 0.559 (1) | T1088 | 0.799 (1) | 0.792 (1) | 0.799 (1) |
| T1047s2 | 0.773 (1) | 0.773 (1) | 0.853 (3) | T1089 | 0.987 (1) | 0.991 (2) | 0.991 (2) |
| T1048 | 0.813 (1) | 0.813 (1) | 0.813 (1) | T1090 | 0.915 (1) | 0.915 (1) | 0.930 (3) |
| T1049 | 0.934 (1) | 0.938 (3) | 0.938 (3) | T1091 | 0.831 (1) | 0.904 (1) | 0.904 (2) |
| T1050 | 0.977 (1) | 0.977 (1) | 0.977 (1) | T1092 | 0.922 (1) | 0.918 (3) | 0.930 (3) |
| T1052 | 0.687 (1) | 0.697 (3) | 0.697 (3) | T1093 | 0.936 (1) | 0.936 (4) | 0.936 (1) |
| T1053 | 0.975 (1) | 0.976 (1) | 0.981 (3) | T1094 | 0.910 (1) | 0.915 (5) | 0.921 (4) |
| T1054 | 0.923 (1) | 0.924 (3) | 0.924 (3) | T1095 | 0.940 (1) | 0.940 (4) | 0.942 (2) |
| T1055 | 0.898 (1) | 0.905 (3) | 0.905 (3) | T1096 | 0.559 (1) | 0.570 (3) | 0.570 (3) |
| T1056 | 0.969 (1) | 0.983 (1) | 0.983 (2) | T1098 | 0.719 (1) | 0.719 (1) | 0.720 (2) |
| T1057 | 0.955 (1) | 0.954 (5) | 0.955 (1) | T1099 | 0.866 (1) | 0.857 (5) | 0.872 (2) |
| T1058 | 0.956 (1) | 0.960 (3) | 0.960 (3) | T1100 | 0.927 (1) | 0.913 (5) | 0.941 (2) |
| T1060s2 | 0.926 (1) | 0.926 (1) | 0.926 (1) | T1101 | 0.934 (1) | 0.937 (2) | 0.938 (3) |
